# Single immunization with recombinant ACAM2000 vaccinia viruses expressing the spike and the nucleocapsid proteins protect hamsters against SARS-CoV-2 caused clinical disease

**DOI:** 10.1101/2021.12.02.470987

**Authors:** Yvon Deschambault, Jessie Lynch, Bryce Warner, Kevin Tierney, Denise Huynh, Robert Vendramelli, Nikesh Tailor, Kathy Frost, Stephanie Booth, Babu Sajesh, Kyle LeBlanc, Christine Layne, Lisa Lin, Daniel Beniac, Michael Carpenter, David Safronetz, Xuguang Li, Darwyn Kobasa, Jingxin Cao

## Abstract

Increasing cases of SARS-CoV-2 breakthrough infections from immunization with predominantly spike protein based COVID-19 vaccines highlight the need for alternative vaccines using different platforms and/or antigens. In this study, we expressed SARS-CoV-2 spike and nucleocapsid proteins in a novel vaccinia virus ACAM2000 platform (rACAM2000). Following a single intramuscular immunization, the rACAM2000 co-expressing the spike and nucleocapsid proteins induced significantly improved protection against SARS-CoV-2 challenge in comparison to rACAM2000 expressing the individual proteins in a hamster model, as shown by reduced weight loss and quicker recovery time. The protection was associated with reduced viral loads, increased neutralizing antibody titre and reduced neutrophil-to-lymphocyte ratio. Thus, our study demonstrates that the rACAM2000 expressing a combination of the spike and nucleocapsid antigens is a promising COVID-19 vaccine candidate and further studies will investigate if the rACAM2000 vaccine candidate can induce a long lasting immunity against infection of SARS-CoV-2 variants of concern.

## INTRODUCTION

The impact of coronavirus disease 2019 (COVID-19), caused by severe acute respiratory syndrome coronavirus 2 (SARS-CoV-2), on global public health and socioeconomic status is unprecedented. To control the COVID-19 pandemic, effective and safe vaccines are essential. Since the start of the pandemic at the end of 2019, there have been almost 300 COVID-19 vaccine candidates that are in clinical or pre-clinical development^1^. To date, COVID-19 vaccines based on several different platforms have been approved for human use, including mRNA vaccines (Pfizer/BNT162b2, Moderna/mRNA-1273), adenovirus vector based vaccines (Oxford/AstraZeneca, Janssen-Johnson & Johnson, Sputnik V) and inactivated virus vaccines (SinoVac, Covaxin). It is an extraordinary achievement that COVID-19 vaccines have been successfully deployed for clinical application in humans within one year after the onset of the pandemic. Clinical data have shown that a nationwide mass immunization with Pfizer/BNT162b2 mRNA vaccine could effectively curb the spread of the disease^2, 3^. Thus, it is encouraging that the COVID-19 pandemic can be contained with a vaccination approach. However, in consideration of the continuous spread of the virus and increasing cases of breakthrough infections from the immunization with current vaccines^4–8^, new vaccines developed with alternative platforms and/or antigens that are safe, induce effective and long-lasting protection against emerging breakthrough variants, and are also convenient for delivery are still needed.

Viral vectors, including vaccinia virus (VACV), are being used as major platforms for development of COVID-19 vaccine candidates^1^. VACV, best known for its role as the vaccine for the eradication of smallpox, has been widely used as a vector for development of various recombinant vaccines^9^. Due to its safety record, the highly attenuated VACV strain, modified vaccinia Ankara (MVA), has been the most widely used vaccinia vector^10^. Currently, several MVA based COVID-19 vaccine candidates have been reported^11–17^. VACV ACAM2000 is another FDA-licensed smallpox vaccine and was derived by plaque-purification from the VACV NYCBH strain^18^, one of the main vaccines used in the eradication of smallpox. Although more immunogenic than MVA, ACAM2000 has not been used as a vector for development of recombinant vaccines due to its adverse effects in humans^19, 20^.

VACV E3L and K3L genes encode two potent inhibitors of type-1 interferon-induced antiviral pathways^21, 22^. Previously it has been shown that deletion of VACV E3L or K3L gene would render the deletion mutant viruses (VACVΔE3L or VACVΔK3L) attenuated^23, 24^. In addition, it has been shown that VACVΔE3L was more potent at inducing innate immune responses than the parental VACV^25, 26^. Thus, it is tempting to use VACVΔE3L as a vector to develop recombinant vaccines^27^. However, VACVΔE3L has a restricted host range and the capacity of VACVΔE3L at delivering antigens is compromised in non-permissive host cells, such as human cells^28^. Recently, we reported that the host range of VACVΔE3L can be modified by swapping VACV K3L gene with a poxvirus K3L orthologue, such as taterapox virus 037 (TATV037), which would enable the virus replication in human cells^29^. Based on the host range specificity of poxvirus K3L genes, we have developed an easy and highly efficient method to construct recombinant VACV using the E3L and K3L double deletion mutant (VACVΔE3LΔK3L) as the backbone^30^. Using a laboratory VACV strain Western Reserve (WR), we recently constructed a recombinant VACVΔE3L expressing the respiratory syncytial virus (RSV) F protein, in that its original K3L gene was replaced with a poxvirus K3L orthologue, TATV037. We demonstrated that the VACVΔE3L vectored recombinant virus was as immunogenic as MVA vector expressing RSV F protein to induce protection against RSV challenge in cotton rats. Importantly, the recombinant virus was highly attenuated in mice^31^.

In this study, we constructed a VACVΔE3LΔK3L based on the FDA approved VACV strain ACAM2000 as a novel recombinant vector for the development of COVID-19 vaccines. In comparison with the MVA platform, recombinants constructed based on this ACAM2000ΔE3LΔK3L backbone have two novel features. First, the E3L gene, which encodes a potent inhibitor of innate immune responses (e.g. IFN and TNF-α pathways)^25, 26^, was deleted, thus the recombinants should be more immunogenic. Second, exchange of VACV K3L with the poxvirus ortholog TATV037 renders the virus replication-competent in human cells^32^, thus increasing the capacity of the recombinants to express the candidate antigens. Since most of current COVID-19 vaccines use the spike (S) protein as an immunogen and the escape SARS-CoV-2 variants from the vaccine induced neutralization immune response have been on the rise^33^, a different immunogen is desirable for the future vaccine development. Therefore, we used the ACAM2000ΔE3LΔK3L as a backbone to construct three recombinant COVID-19 vaccine candidates expressing SARS-CoV-2 S (rACAM2000S), nucleocapsid (N, rACAM2000N), and a combination of the S and N (rACAM2000SN). A poxvirus K3 orthologue TATV037, which will enable ACAM2000ΔE3LΔK3L replication-competent in human and rodent cells was used as a selection marker^30^. Hamsters were immunized with a single intramuscular (i.m.) injection of the rACAM2000 viruses and the protective efficacy was examined following intranasal (i.n.) challenge with SARS-CoV-2 virus.

## MATERIALS AND METHODS

### Cells and Viruses

The SARS-CoV-2 hCoV-19/Canada/ON-VIDO-01/2020 isolate was described previously^34^. VACV ACAM2000 was a gift from Dr. David Evans (University of Alberta, Canada). BHK21, HeLa and Vero cells were from ATCC, and A549/PKR-RNaseL-cells were from Dr. B. Moss (NIH, USA)^35^.

### Construction of recombinant ACAM2000 VACV

The E3L and K3L genes of the ACAM2000 VACV were deleted with the method similar to what we previously described with VACV WR strain using the standard homologous recombination procedure for making recombinant poxviruses with the following modifications^32^. The detailed information is outlined in supplemental Figure 1. First, the E3L gene was disrupted using a transient selection method similar to the previously described^36^, except an mCherry fluorescent protein was used as the transient selection marker (supplemental Figure 1A). The purified E3L deletion virus (ACAM2000ΔE3L) does not carry the selection marker mCherry, as colorless plaques were selected during the plaque purification. Then, the K3L gene was disrupted by insertion of an EGFP gene driven by the VACV late promoter p11 using the standard homologous recombination procedure^37^ (supplemental Figure 1B). The ACAM2000ΔE3L with the disrupted K3L gene (ACAM2000ΔE3LΔK3L) was selected and purified using EGFP as the selection marker. A549/PKR-RNaseL-cells were used to select both ACAM2000ΔE3L and ACAM2000ΔE3LΔK3L viruses. The purified ACAM2000ΔE3LΔK3L was confirmed with PCR and sequencing of the respective genomic loci (data not shown).

Previously, we demonstrated that replacement of the K3 gene of VACVΔE3L with the taterapox virus K3 ortholog TATV037 could render the virus replication-competent in both hamster BHK21 and human HeLa cells^32^. Using a poxvirus K3 ortholog gene as a selection marker, we have developed a novel method to construct recombinant vaccinia viruses^30^. Based on this protocol, we used the ACAM2000ΔE3LΔK3L as the backbone and the TATV037 as a selection marker to construct recombinant ACAM2000 (rACAM2000) expressing the wild-type full-length SARS-CoV-2 S and N proteins separately or in combination. The genes were synthesized by Genscript (NJ, USA) based on the Wuhan isolate with a FLAG tag included at the C-terminus (Genbank accession number MN908947). Silent mutations were made to remove the poxvirus early transcription termination signal, TTTTTNT, in the genes. The structure of the recombinant shuttle vector is shown in Figure 1A. The TATV037 gene is highly homologous with VACV K3L^32^ and the TATV037 gene was used as one of the flanking sequence for homologous recombination. Since the native VACV K3L promoter likely resides in the adjacent K4L gene, which forms a part of the flanking sequence, the promoter for a sheeppox virus K3L ortholog gene was used to drive the transcription of TATV037 to avoid unwanted in-genome homologous recombination. SARS-CoV-2 S and N genes were driven by a vaccinia early and late promoter, mH5^38^. The A549/PKR-RNase L-cells infected with ACAM2000ΔE3LΔK3L virus expressing EGFP were transfected with the respective recombination shuttle vectors. The recombinant virus, which does not express EGFP, were selected and purified in BHK21 cells. Expression of the S and N proteins were confirmed by Western blotting with a FLAG antibody. All the rACAM2000 recombinant stocks used for animal experiments were partially purified through a 36% sucrose cushion^39^.

**Figure 1.**
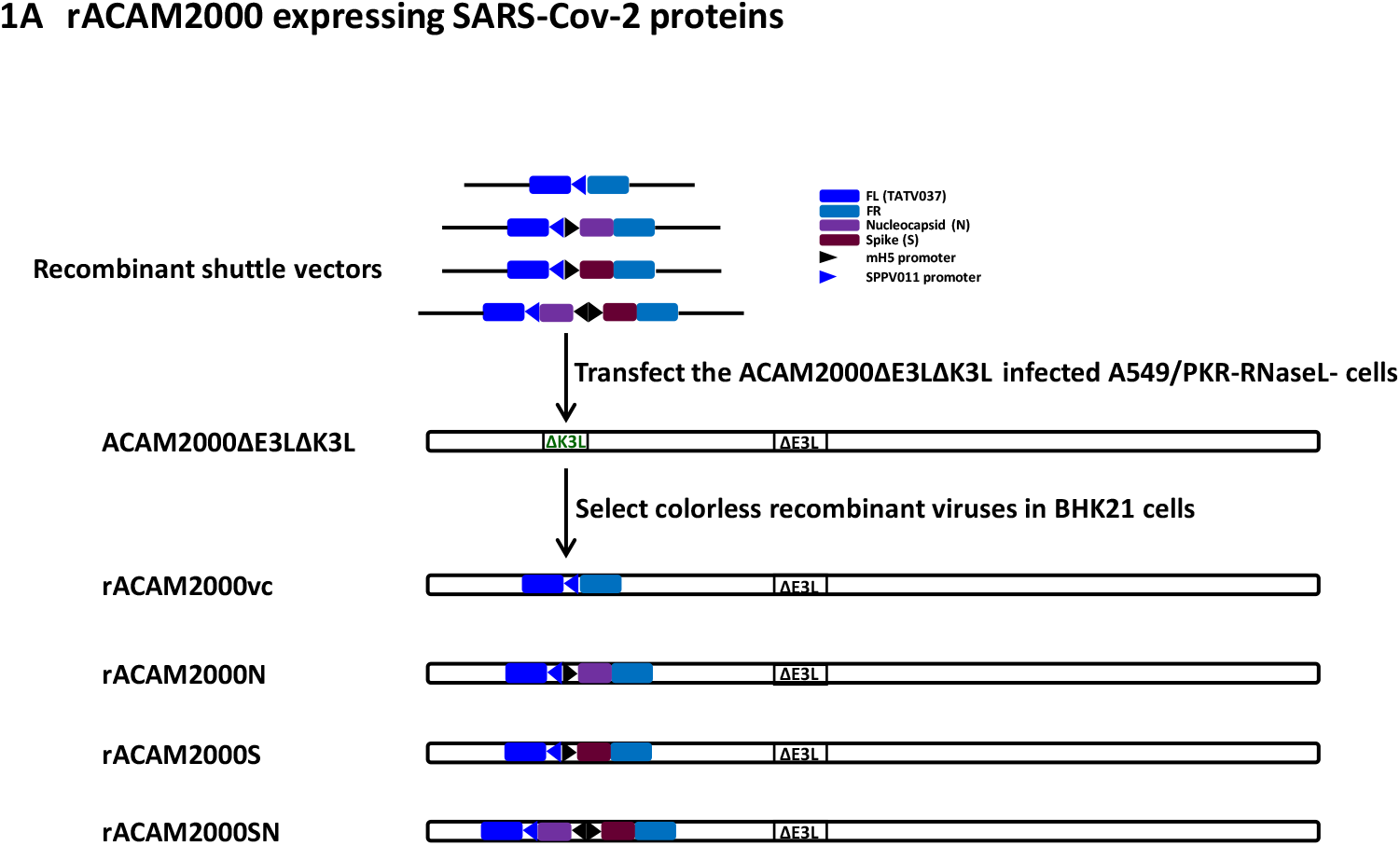
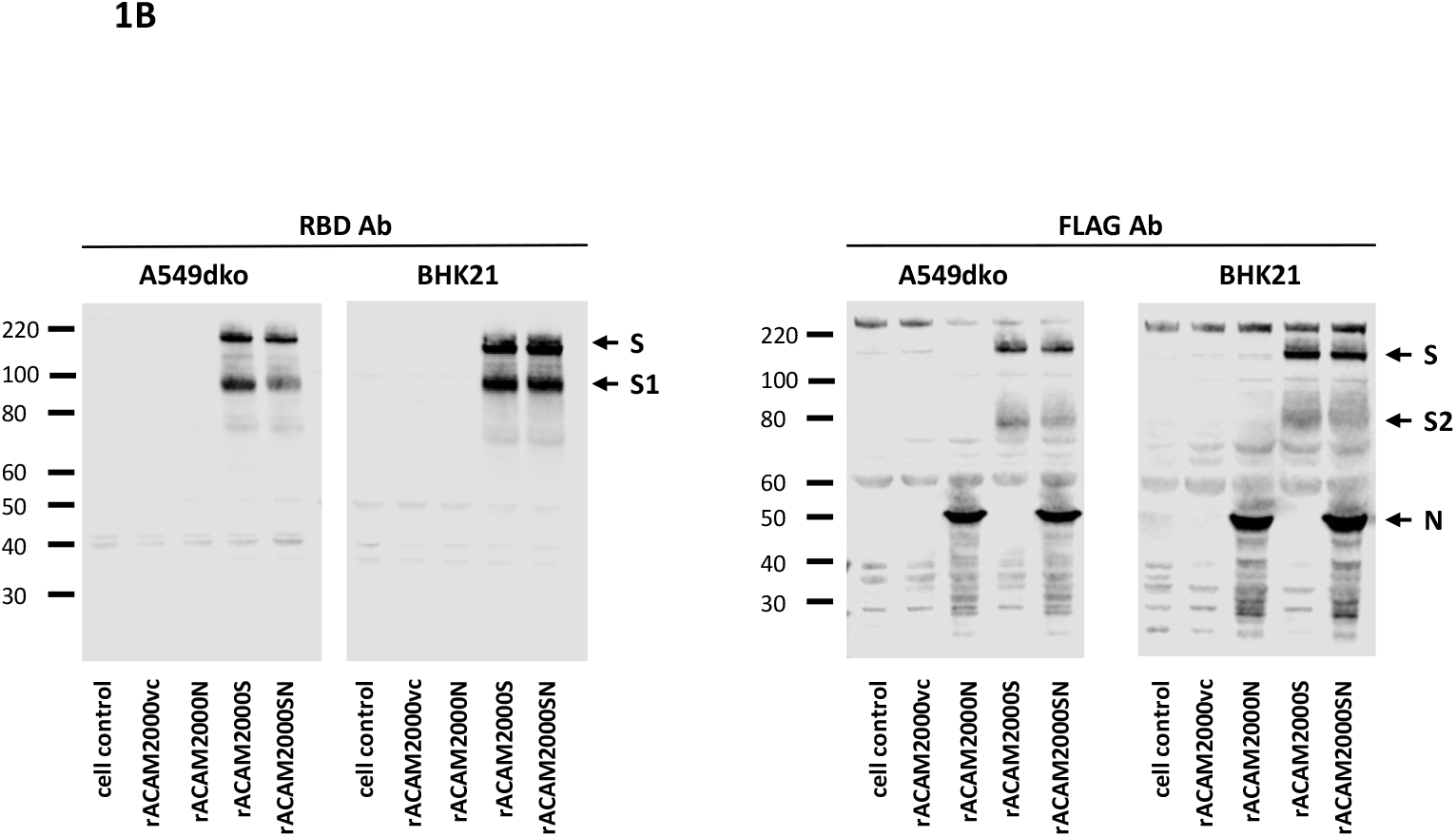
Construction of recombinant ACAM2000 (rACAM2000) expressing SARS-CoV-2 spike (S) and nucleocapsid (N) proteins. A: Schematic illustration of construction of recombinant ACAM2000ΔE3LΔK3L expressing SARS-CoV-2 S and N proteins. The shuttle vectors to mediate the integration of the SRAS-Cov-2 S and/or N genes into the K3L locus consist of a K3L ortholog gene, taterapox virus 037 (TATV037) as a positive selection marker driven by the promoter of a sheeppox virus K3L ortholog gene SPPV011, and the SARS-CoV-2 S and N genes driven by the vaccinia early and late promoter mH5. The final recombinant ACAM2000ΔE3LΔK3L expressing SARS-CoV-2 spike (S) and nucleocapsid (N) proteins had the EGFP gene replaced by the SARS-CoV-2 genes, and thus have no fluorescence protein marker. Recombinant ACAM2000 virus naming conventions used are as follows: rACAM2000vc: the empty control recombinant not expressing SARS-CoV-2 protein, rACAM2000N: the recombinant expressing SARS-CoV-2 N protein, rACAM2000S: the recombinant expressing SARS-CoV-2 S protein, rACAM2000SN: expressing both S and N proteins. B: Western blot analysis of SARS-CoV-2 S and N proteins. Antibodies to the RBD were used to detect the full-length S and S1 and FLAG antibody was used to detect the N, the full-length S and S2 proteins. Expression of the proteins were examined in BHK21 and A549dko cells.

### Western blot analysis

HeLa and BHK21 Cell monolayers in a 12-well plate were infected with rACAM2000 viruses at an moi of 5 and the cell lysate was collected at 16 hours post infection with 200 µl of protein sample buffer. SDS-PAGE, membrane transfer, antibody blotting, and protein bands detection were performed as described previously^32^. The antibody for SARS-CoV-2 S RBD was from Sino Biological Inc (cat # 40592-T62-100), and the antibody for the FLAG tag was from Sigma (cat # F7425).

### Animal experiment

Five to seven weeks old Golden Syrian hamsters were purchased from Charles River Laboratories (USA), and males and females were caged separately. All the procedures were approved by the Animal Care Committee of the Canadian Science Centre for Human and Animal Health and followed the guidelines of the Canadian Council for Animal Care.

For the immunization, 2×10^7^ PFU of the rACAM2000 recombinants in 200 µl of serum free DMEM medium were administered through i.m. injection in the right and left quadriceps muscles in a BSL2 lab. For immunization with the combined rACAM2000S and rACAM2000N, the two recombinants were injected separately in the right and left quadriceps muscles. For the SARS-CoV-2 challenge, 10^5^ TCID_50_ of the virus in 100 µl serum free DMEM was administered through the i.n. route (50 µl per nostril) in the biosafety level 4 laboratory at the National Microbiology Laboratory.

The workflow of the hamster experiment is illustrated in Figure 3. Prior to the start of the experiment, serum samples were collected from all hamsters. 12 male and 12 female animals were immunized with each of the rACAM2000 recombinants (Figure 2) through i.m. injection and the weight of hamsters was monitored daily for one week post immunization. Three weeks post immunization, serum samples were collected and hamsters were challenged with 10^5^ TCID_50_ of SARS-CoV-2 through the i.n. route and weight change and other clinical signs were monitored daily. On day 5 post challenge, 6 male and 6 female animals immunized with each rACAM2000 were euthanized. Blood and various tissues were collected for analysis. The remaining 6 male and 6 female hamsters of each group were monitored up to 28 days post challenge and then euthanized.

**Figure 2.**
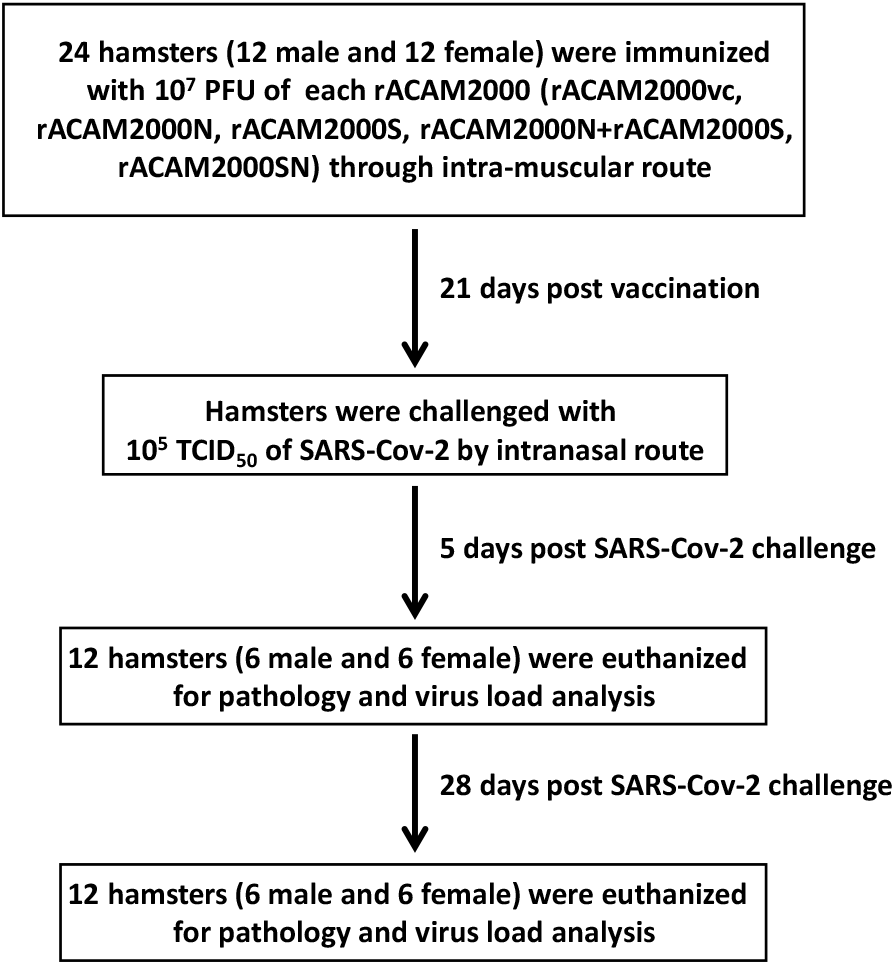
Timeline of the vaccination and SARS-CoV-2 challenge.

### Determination of infectious viral titres by TCID_50_ assay

Nasal turbinate, trachea, lung, small intestine and liver tissues were weighed and homogenized with stainless steel beads in 1 ml of MEM (1% FCS and 1% Pen/Strep) for 30 seconds at 4 m/sec on the Bead Ruptor Elite (Omni International). The tissue debris was removed by centrifugation at 1500 x g for 10 minutes. 100 µl of a series of 10 fold-dilutions of the tissue supernatant was added in triplicate to Vero cell monolayers in a 96-well plate. Following incubation at 37°C, and 5% CO_2_ for 5 days, the cytopathic effect (CPE) was visualized under a microscope and the TCID_50_ per gram of tissue was calculated using the Reed-Muench method^40^.

### Molecular determination of the viral load by quantitative real-time PCR (qRT-PCR)

Total RNA was extracted from hamster tissue samples using the RNEasy Mini kit (Qiagen) according to the manufacturer’s instruction. The qRT-PCR was performed using TaqPath 1-Step Multiplex Master Mix (Applied Biosystems) and primer/probe sets against SARS-CoV-2 N gene (For: GAC CCC AAA ATC AGC GAA AT; Rev: TCT GGT TAC TGC CAG TTG AAT CTG; probe: ACC CCG CAT TAC GTT TGG TGG ACC) and E gene (For: ACA GGT ACG TTA ATA GTT AAT AGC GT; Rev: ATA TTG CAG CAG TAC GCA CAC A; probe: ACACTA GCC ATC CTT ACT GCG CTT CG) (IDT). Reactions were performed in MicroAmp Fast Optical 96-well plates (Applied Biosystems) on a StepOnePlus real time PCR machine (Applied Biosystems). The qRT-PCR cycle consisted of an initial step of 53°C for 10 minutes, followed by 95°C for 2 minutes, followed by 40 cycles of 95°C for 2 seconds and 60°C for 30 seconds. Viral load was determined using standard curve generated with a serially diluted synthetic standard (BEI Resources).

### Microneutralization assay

The microneutralization assay was performed similar to the recently described protocol^41^. Briefly, hamster sera were heat-inactivated at 56°C for 1 hour. 100 µl of a series of 10-fold dilutions of the heated-inactivated sera were mixed with 100 plaque forming units (PFU) of SARS-CoV-2 and was incubated at 37°C for 1 hour. Triplicates of the serum and virus mixture were added to Vero cells in 96-well plates. The CPE was visualized at 5 days post infection. The highest dilution of the sera, at which 50% of the infectivity was neutralized, was determined as NT_50_ using the Reed-Muench formula^40^.

### ELISA

MaxiSorp 96-well flat bottom immuno-plates (ThermoFisher) were coated with 10 μg/mL of the full-length SARS-CoV-2 N or S protein based on Wuhan strain of SARS-CoV-2 at 4°C overnight. The highly purified S and N proteins were prepared at the National Microbiology Laboratory by Dr. M. Carpenter using a pcDNA3.1/HEK293 eukaryotic expression system (unpublished). The following day, the plates were washed 3 times with 200µl PBS supplemented with 0.1% Tween-20 (PBST). After the final wash, plates were blocked with 100µl ChonBlock Blocking/Sample Dilution buffer (Chondrex) at room temperature for 1 hour. Blocking buffer was then removed and plates were washed 3 times in 200µl PBST. Hamster sera samples were diluted with ChonBlock buffer to 100µl and plates were incubated at room temperature for 2 hours. Then the hamster sera was removed and plates were washed 5 times 300µl with PBST. A 1:1000 dilution of Goat anti-hamster IgG HRP conjugated antibody (ThermoFisher) was prepared in ChonBlock Secondary Antibody Dilution Buffer (Chondrex), 100µl was added and plates were incubated at room temperature for 1 hour. Secondary antibody was removed and plates were washed 5 times in 300µl PBST. Plates were incubated with 75µl KPL SureBlue Reserve TMB Microwell Peroxidase Substrate (Seracare) at room temperature for 10 minutes. Reaction was stopped using an equal volume of 1N H_2_SO_4_. Plates were read at 450nm on a SpectraMax Plus spectrophotometer (Molecular Devices).

### Serum Biochemistry and Hematology

Serum biochemistry and hematological analyses were performed using Abaxis HM5 and Abaxis VetScan VS2 respectively (Abaxis Veterinary Diagnostics) as previously described^34^.

### Histopathology

Excised tissues of various animal organs were fixed in 10% neutral buffered formalin solution, and processed for paraffin embedding. Paraffin blocks were cut into 5-µm-thick sections that were then mounted on Shandon™ positively charged glass slides. Sectioned tissues were rehydrated through 3 changes of xylene for 3 minutes followed by 2 changes of 100% and 95% ethanol for 90 seconds. This was followed by a 90 second water rinse and then 2 changes of hematoxylin for 90 seconds, 90 seconds bluing solution and 90 seconds decolorizer, 90 seconds Eosin. Tissues were then dehydrated with 95% ethanol for 90 seconds, followed by 2X changes of 90 seconds of 100% ethanol. The final treatment with xylene for 5 minutes was used to remove paraffin. Slides were then cover slipped with Permount™ mounting medium. All slides were imaged with ZEISS Axio Scan.Z1 slide scanner.

Semi-quantitative visual image analysis was performed using a scoring system to estimate the extent of interstitial pneumonia across each section. Each lung lobe was analyzed and the level of involvement estimated as a percentage across the slide in comparison to non-infected control hamsters.

### Statistic analysis

Data were analyzed and plotted using GraphPad Prism 9 software. The significance of the difference between two groups of hamsters was analyzed using a two-tailed nonparametric Mann-Whitney u test. A P value greater than 0.1 was considered not significant.

## RESULTS

### Expression of SARS-CoV-2 S and N proteins

As shown in Figure 1A, the SARS-CoV-2 S and N genes driven by a VACV early and late promoter mH5 were inserted at the K3 locus in the genome of VACV ACAM2000ΔE3LΔK3L using a similar strategy as previously described^30, 31^. Human cells (A549) and hamster cells (BHK21) were infected with the rACAM2000 recombinants and expression of the S and N proteins were examined with Western blot analysis using an antibody against the RBD region of the S and an antibody for the FLAG tag added to the C-terminus of the proteins. As shown in Figure 1B, the RBD antibody detected the full-length S and S1 from the cell lysates infected with the rACAM2000S and rACAM2000SN recombinants, while the FLAG antibody detected the full-length S, S2 and the N proteins in the samples from corresponding rACAM2000 recombinant infected cells.

### The rACAM2000 expressing SARS-CoV-2 S and N proteins protect hamsters from weight loss following SARS-CoV-2 challenge

It has been shown that NYCBH strain VACVΔE3L, from which the ACAM2000 VACV was derived, were highly attenuated in mice^23^. Since VACVΔE3L replicates equally well in both mouse and hamster cells^29^, it is expected that the virus would be attenuated in hamsters. Nonetheless, the weight change and visible clinical signs of all animals were monitored following the i.m. vaccination with the rACAM2000 recombinant viruses shown in Figure 1A. Body weight was recorded daily for 7 days post vaccination and the percentage of weight change is shown supplemental Figure 2. Most of the animals gained weight at 7 days post vaccination. Although a few hamsters vaccinated with rACAM2000N, rACAM2000S or rACAM2000SN lost a moderate amount of weight in the first week post vaccination, their body weight recovered prior to the challenge with SARS-CoV-2 at day 21 post vaccination (data not shown). No other visible clinical signs were observed before the SARS2-Cov2 challenge. A similar observation was also made in mice from our recent study with the VACVΔE3LΔK3L WR strain expressing a RSV F protein^31^.

Three weeks post vaccination, hamsters were challenged with 10^5^ TCID_50_ of SARS-CoV-2 through the i.n. route. On day 5 post challenge, 12 animals (6 males and 6 females) of each vaccinated group were euthanized, underwent necropsy, and tissues were collected for histopathological examination and viral load analysis. Another 12 hamsters (6 males and 6 females) were monitored for up to 28 days and body weights were recorded daily for 15 days post challenge. Following SARS-CoV-2 challenge, the major clinical signs shown by hamsters were weight loss and lethargy. Since weight loss is quantifiable, the percentage of weight change was used as the main indicator of clinical disease. The daily weight changes of each hamster is shown in Figure 3A. The percentage of the maximum weight change and the DPI (day post infection) when the maximum weight loss occurred are summarized in Figure 3B and 3C respectively. In the rACAM2000S vaccinated group, the maximum weight loss occurred 1 day earlier (5 DPI vs 6 DPI, P<0.1) than the rACAM2000vc group on average, while the difference in the maximum weight loss was not statistically significant (9.5% vs 12.2% in comparison with the rACAM2000vc). The rACAM2000N induced moderate but statistically significant protection as measured by maximum weight loss and the DPI when maximum weight loss occurred (7.9% vs 12.2% in the vector control, 5DPI vs 6 DPI of the vector control, P<0.01 and P<0.1 respectively, Figure 3B and 3C). In comparison to the rACAM2000vc, rACAM2000S and rACAM2000N groups, the maximum weight loss of the hamsters vaccinated with rACAM2000S plus rACAM2000N or rACAM2000SN (expressing S and N from the same recombinant virus) were significantly less (5% and 5.4% vs 12.2% in the rACAM2000vc group, P<0.0001, Figure 3B). In addition, the maximum weight loss occurred markedly earlier following the vaccination with the recombinants expressing S and N antigens than other groups (a median of 3 or 4 DPI vs 5 or 6DPI, P<0.0001 and P<0.001 respectively, Figure 3C).

**Figure 3.**
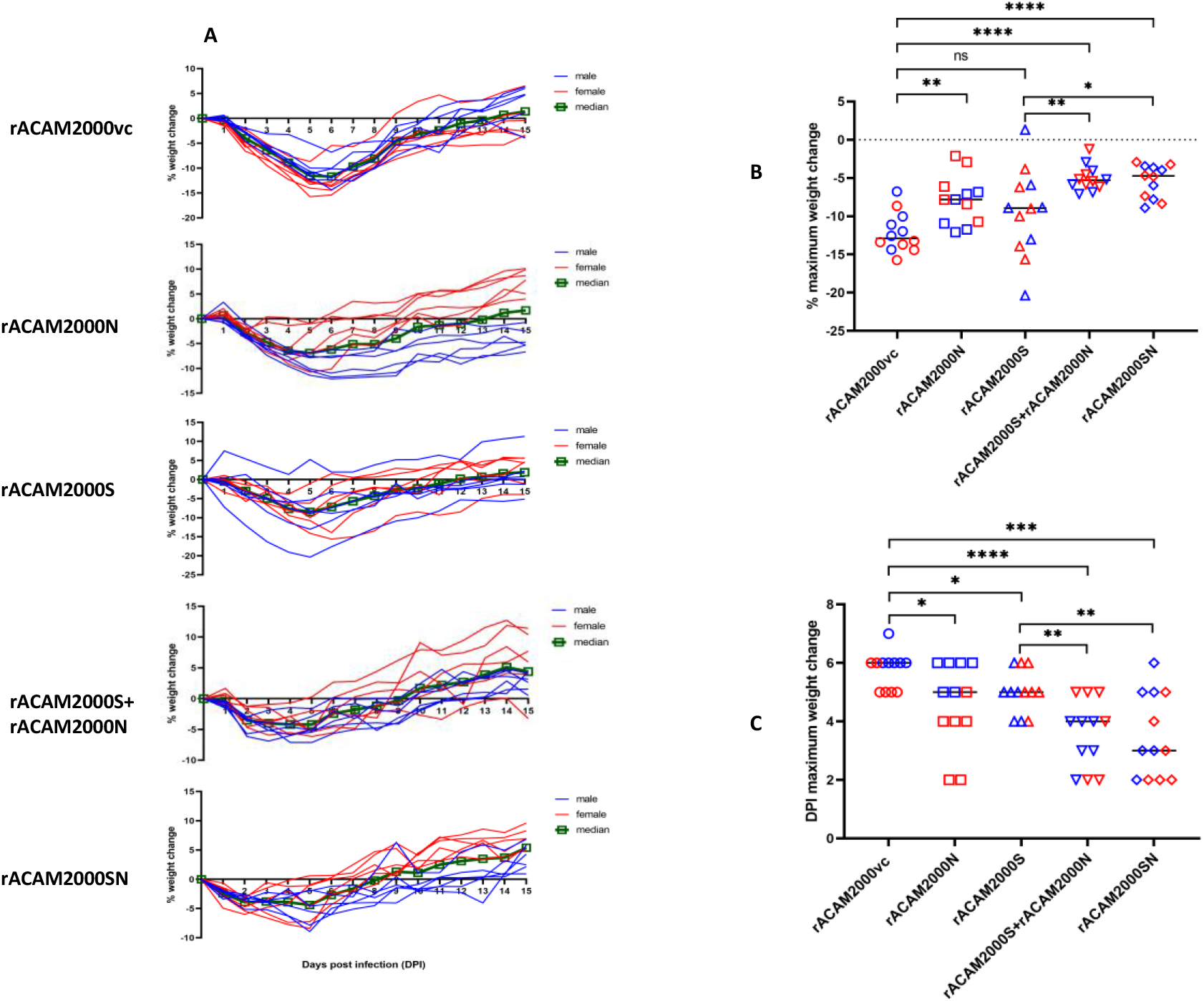
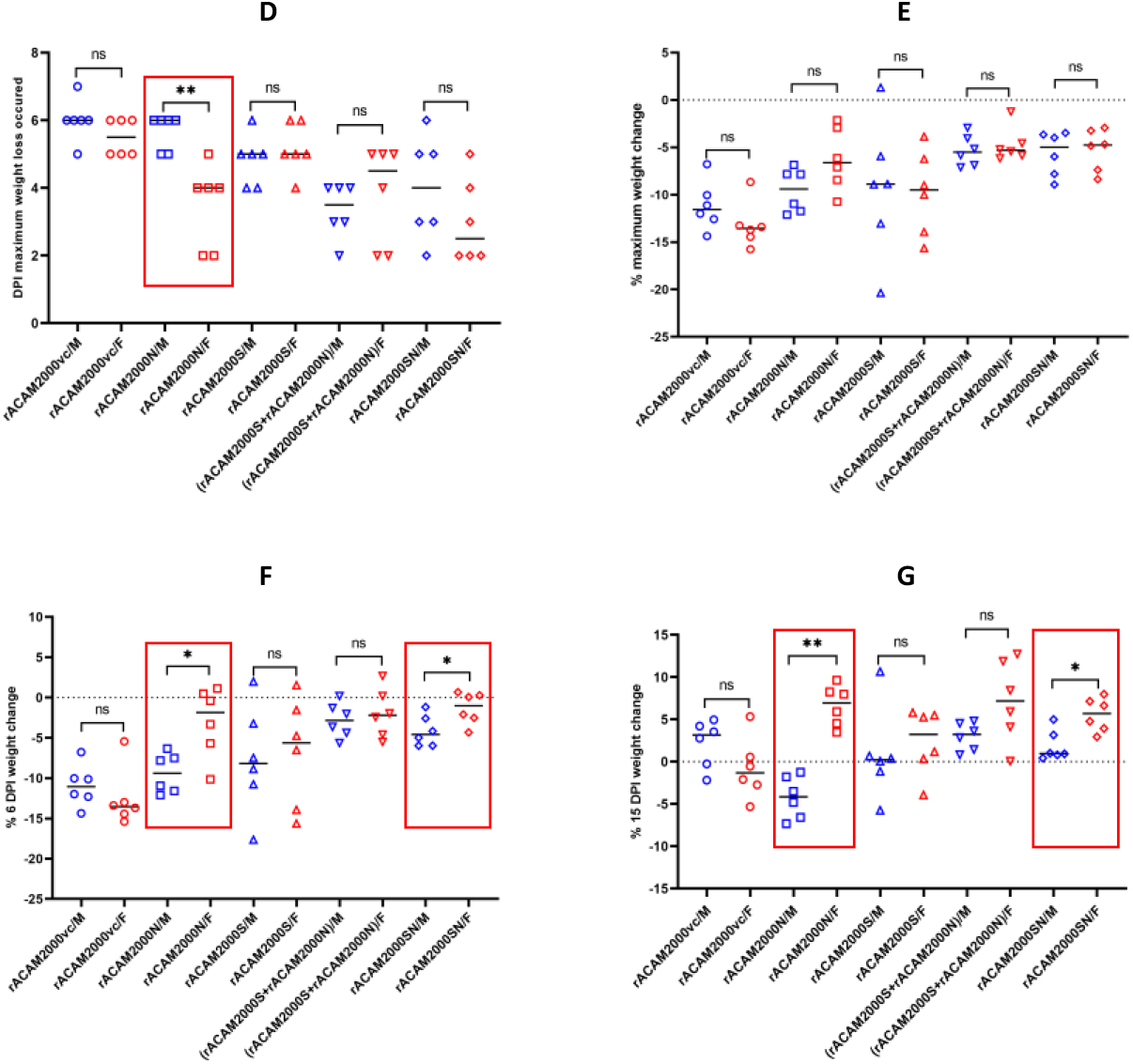
Weight loss of hamsters following SARS-CoV-2 challenge. A. Percentage of daily weight change of individual hamsters up to 15 days post SRAS-Cov-2 challenge. Blue line represents male hamsters, red line represents female hamsters, and median weight change is shown in green. B. The maximum weight loss post SARS-CoV-2 challenge. C. The day post infection (DPI) when the maximum weight loss observed. D. Comparison of the DPI the maximum weight loss occurred between male and female hamsters. E. Comparison of the maximum weight loss between male and female hamsters. F. Comparison of the weight changes at 6 DPI between male and female hamsters. G. Comparison of the weight changes at 15 DPI between male and female hamsters. The circle shape represents rACAM2000vc, square for rACAM2000N, triangle for rACAM2000S, upside-down triangle for rACAM2000S+rACAM2000N, and diamond for rACAM2000SN; red-colored shape for female and blue-colored shape for male hamsters. The P values were calculated with two-tailed Mann-Whitney u test using Graphpad Prism 8.0 package, ****:p<0.0001; ***:p<0.001; **:p<0.01; *:p<0.1.

In the rACAM2000N group, we noticed an unexpected weight change pattern associated with the sex of hamsters, in that the body weight of female hamsters immunized with the rACAM2000N recovered noticeably faster than the males (Figure 3A). Thus the difference in weight change between male and female hamsters was further compared (Figure D-G). On average, the maximum weight loss occurred in the male hamsters immunized with rACAM2000N was at 6 DPI, same as the rACAM2000vc group, while the maximum weight loss of female hamsters in this group occurred at 4 DPI (P<0.01, Figure 3D). Thus, the female hamsters immunized with rACAM2000N started to recover from SARS-CoV-2 challenge 2 days earlier than the males. Although the difference in the maximum weight losses was not statistically significant (Figure 3E), the body weight of the female hamsters immunized with rACAM2000N recovered significantly better than the males at the 6 and 15 days post challenge (Figure 3F and 3G). A similar difference was also observed between the male and female hamsters vaccinated with rACAM2000SN, although to a lesser degree than the rACAM2000N group (Figure 3F and 3G). No statistically significant differences in the weight change were observed between the males and the females in other groups.

### Vaccination with the rACAM2000 expressing S significantly reduced the infectious viral loads in lung and nasal turbinate

Nasal turbinate, trachea, lung, small intestine and liver tissues collected from the 5 DPI necropsy were analyzed for infectious viral load by TCID_50_ assay (Figure 4). In the lung, the TCID_50_ per gram of tissue from hamsters vaccinated with rACAM2000S, rACAM2000S+rACAM2000N and rACAM2000SN were at least 100 (2 log value for rACAM2000S and rACAM2000S+rACAM2000N) or 1000 fold (3 log value for rACAM2000SN) less than the rACAM2000vc and rACAM2000N groups (P<0.0001, the detection limit of the assay was 2.1 log, Figure 4A). In the nasal turbinate, the TCID_50_ per gram of tissue of all vaccinated animals was lower than the rACAM2000vc group with statistical significance (P value between <0.1 to <0.001). The vaccinations with the recombinants expressing the S (either S only or the combined S and N: rACAM2000S+rACAM2000N or rACAM2000SN) reduced the viral load in the nasal turbinate more than the rACAM2000N group (P<0.01, Figure 4B). No significant difference in the infectious viral load was observed between males and females in all groups examined (supplemental Figure 3). Only low levels of the virus were observed from some trachea specimen and there was no correlation between the control and vaccinated animals (data not shown). No infectious virus was detected from the liver and small intestine specimen collected at 5 DPI (data not shown).

**Figure 4.**
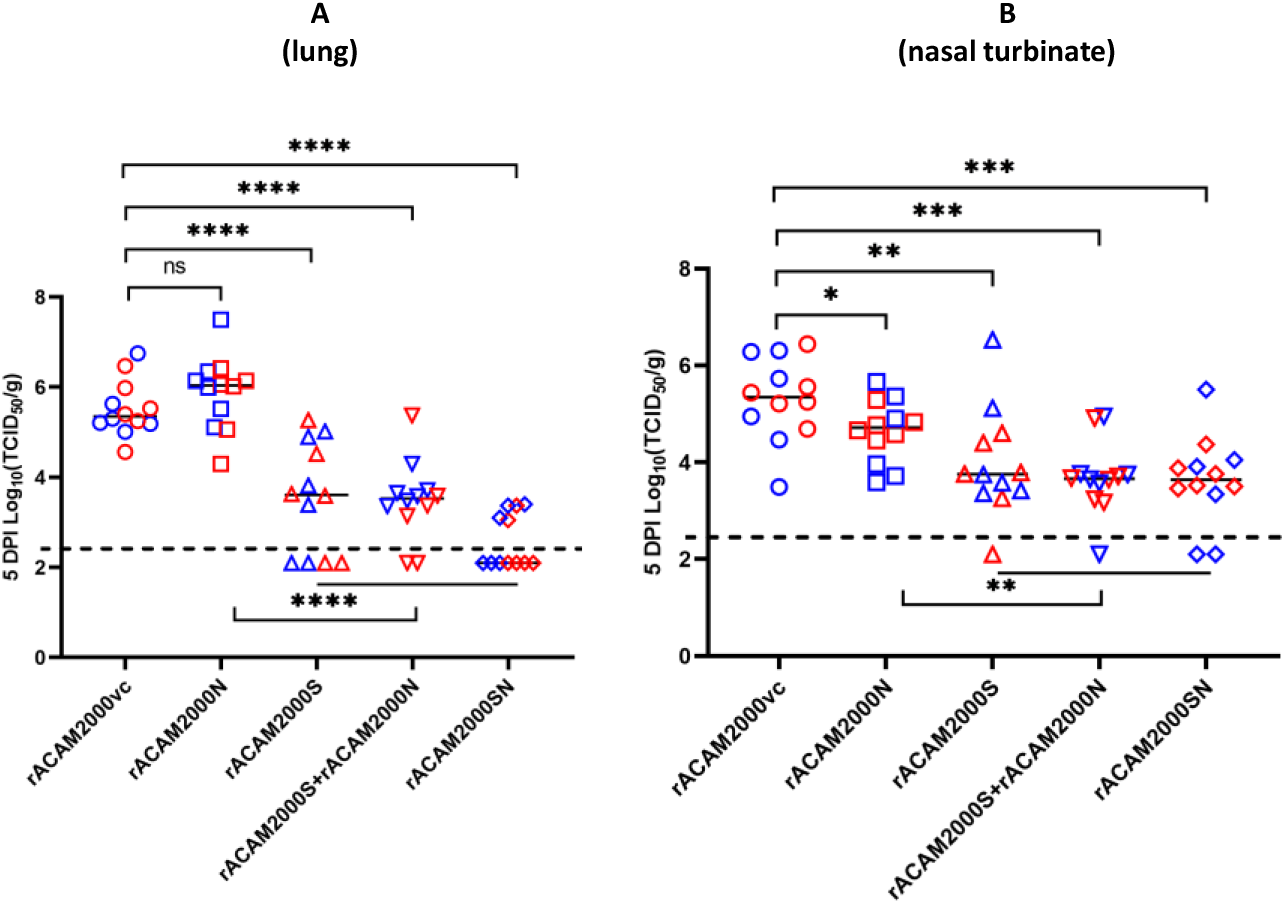
Infectious SARS-CoV-2 viral loads in the lung and nasal turbinate at the 5 DPI. A: Virus titers in the lung determined as reciprocal of the end point dilution TCID_50_ per gram of tissue. The dotted line indicate the limit of detection. B: Virus titers in the nasal turbinate determined the same as in A. The circle shape represents rACAM2000vc, square for rACAM2000N, triangle for rACAM2000S, upside-down triangle for rACAM2000S+rACAM2000N, and diamond for rACAM2000SN; red-colored shape for female and blue-colored shape for male hamsters. The P values were calculated with two-tailed Mann-Whitney u test using Graphpad Prism 8.0 package, ****:p<0.0001; ***:p<0.001; **:p<0.01; *:p<0.1.

### Combined expression of the S and N proteins improved effect on reducing the viral load in distal tissues determined by qRT-PCR

Since the sensitivity of the TCID_50_ assay was limited in detecting the presence of the virus (the limit of detection was 2.1 log value per gram of the tissue), the presence of SARS-CoV-2 in the various tissues was also examined by qRT-PCR. Two primer pairs were employed, data for the SARS-CoV-2 envelope gene is shown in Figure 5 and data for the nucleocapsid gene is shown in supplemental Figure 4. In the respiratory tract tissues (nasal turbinate, trachea and lung), the viral RNA copy numbers were generally higher in the hamsters vaccinated with rACAM2000N than the rACAM2000vc, particularly in the lung and trachea (P<0.1 and 0.01 respectively, Figure 5A, 5C). There were no significant differences between the RNA copy numbers from the nasal turbinate and trachea of rACAM2000S, rACAM2000S+rACAM2000N and rACAM2000SN vaccinated animals in comparison to the rACAM2000vc group. The rACAM2000S+rACAM2000N vaccinated hamsters had lower RNA copies in the lung than the other groups. In the distal tissues, the RNA copy numbers were significantly lower in the hamsters vaccinated with rACAM2000 expressing SARS-CoV-2 antigens (S, N, or S+N) than the rACAM2000vc (statistical significance between P<0.1 to P<0.0001, Figure 5D and 5E). In addition, combining S and N proteins (rACAM2000S+rACAM2000N or rACAM2000SN) in the vaccination reduced the viral loads in the liver and small intestine more than the vaccination with a single SARS-CoV-2 antigen, S or N.

**Figure 5.**
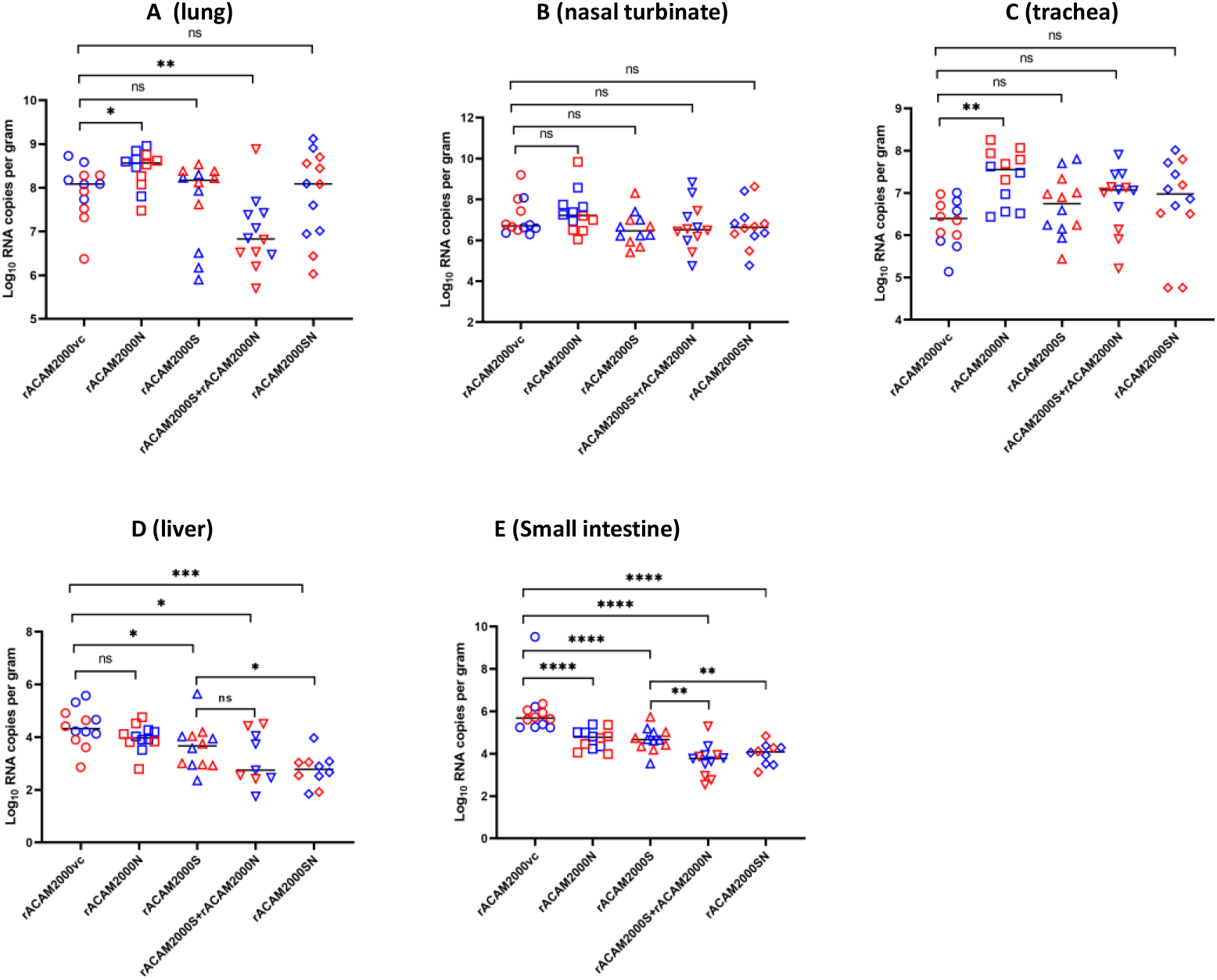
SARS-CoV-2 viral loads in various tissues collected at 5 DPI determined using qRT-PCR specific for the E gene. A: The SARS-CoV-2 RNA copy numbers (per gram of tissue) in the lung. B: The RNA copy numbers in the nasal turbinate. C: The RNA copy numbers in the trachea. D: The RNA copy numbers in the liver. E: The RNA copy numbers in the small intestine. The circle shape represents rACAM2000vc, square for rACAM2000N, triangle for rACAM2000S, upside-down triangle for rACAM2000S+rACAM2000N, and diamond for rACAM2000SN; red-colored shape for female and blue-colored shape for male hamsters. The P values were calculated with two-tailed Mann-Whitney u test using Graphpad Prism 8.0 package, ****:p<0.0001; ***:p<0.001; **:p<0.01; *:p<0.1.

### Antibody responses

Antibody responses were analysed using microneutralization assay with infectious SARS-CoV-2 virus and an ELISA with purified S and N proteins. No antibodies were detected in the sera collected prior to vaccination (data not shown).

Twenty-one days post vaccination, antibodies against S or N protein were detected in the sera from all hamsters vaccinated with the rACAM2000 expressing the corresponding proteins in ELISA (Figure 6A and 6B). However, none of the animals developed detectable neutralizing antibodies prior to SARS-CoV-2 challenge (Figure 6C). On day 5 post challenge, antibody levels significantly increased in both S and N ELISA assays in comparison to the pre-challenge and rACAM2000vc samples (Figure 6A and 6B). Interestingly, S antibody levels in hamsters vaccinated with rACAM2000N was significantly lower than the rACAM2000vc following SARS-CoV-2 challenge (Figure 6A). Additionally, hamsters vaccinated with rACAM2000SN showed lower S antibody level than hamsters immunized with rACAM2000S and rACAM2000S+rACAM2000N at both 21 days post vaccination and 5 days post SARS-CoV-2 challenge, albeit higher than the rACAM2000vc (Figure 6A). Neutralizing antibodies were detected in all hamsters vaccinated with rACAM2000 expressing the S protein (rACAM2000S, rACAM2000SN, and rACAM2000S+rACAM2000N) at 5 days post SARS-CoV-2 challenge with the median logNT_50_ titre between 2.3 and 2.5 (Figure 6C). Following SARS-CoV-2 challenge, 10 of 12 hamsters immunized with the empty control recombinant, rACAM2000vc, developed detectable neutralizing activity with a median logNT_50_ titre of 1.9, which was significantly lower than the hamsters vaccinated with the rACAM2000 expressing the S antigen (P<0.001 for rACAM2000SN, P<0.0001 for rACAM2000S and rACAM2000s+rACAM2000N). Interestingly, only 5 (1 male and 4 females) out of 12 hamsters vaccinated with rACAM2000N developed detectable neutralizing antibody following SAR-CoV-2 challenge, significantly lower than the rACAM2000vc (P<0.01). The median NT_50_ of the rACAM2000SN sera was also lower than the rACAM2000S and rACAM2000S+rACAM2000N, albeit not statistically significant (Figure 6C). For statistical analysis, the NT_50_ titre of those with undetectable neutralizing activity was arbitrarily assigned a value of 1.77, which is the limit of detection and this would not arbitrarily increase the statistical significance of the difference.

**Figure 6.**
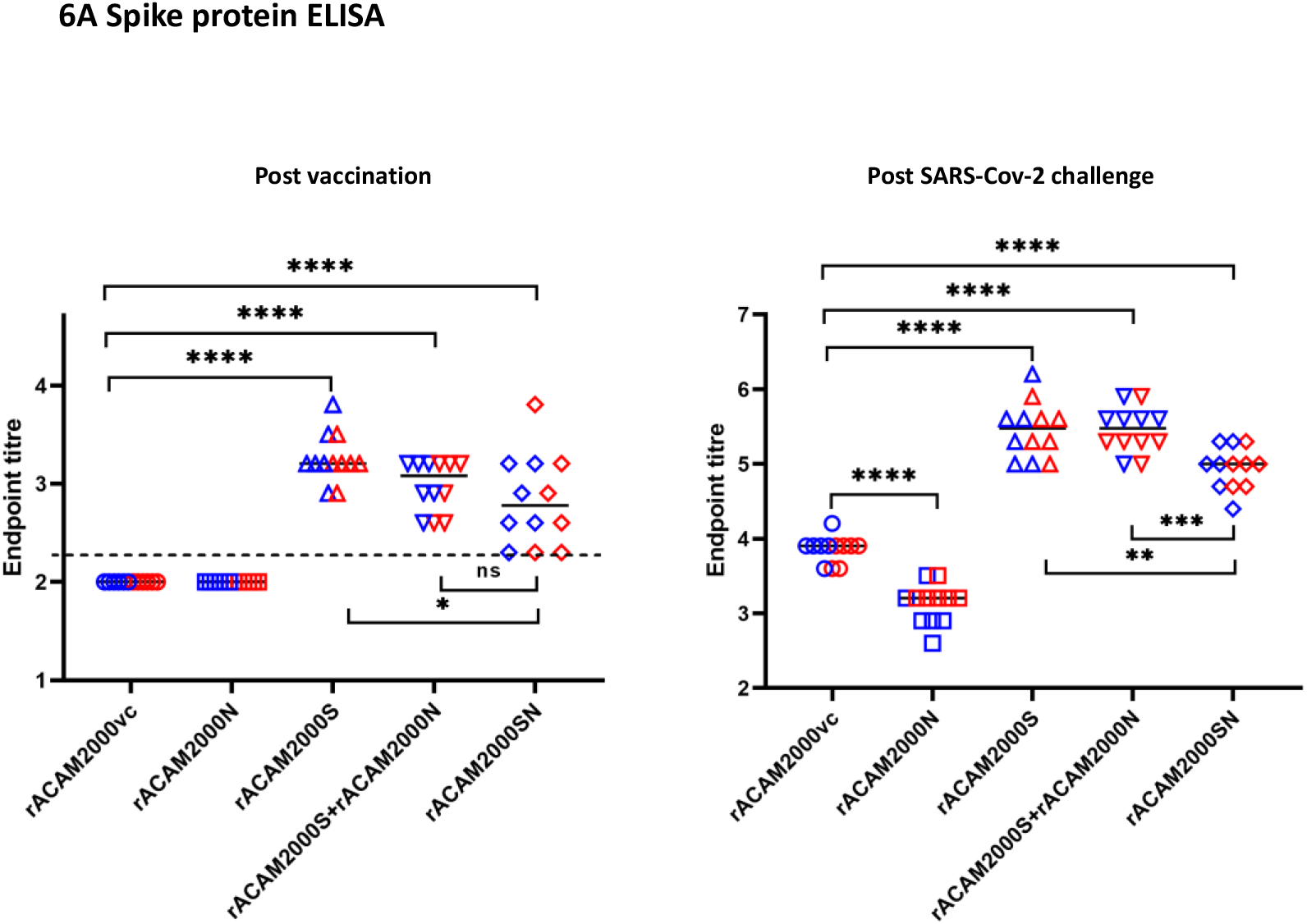
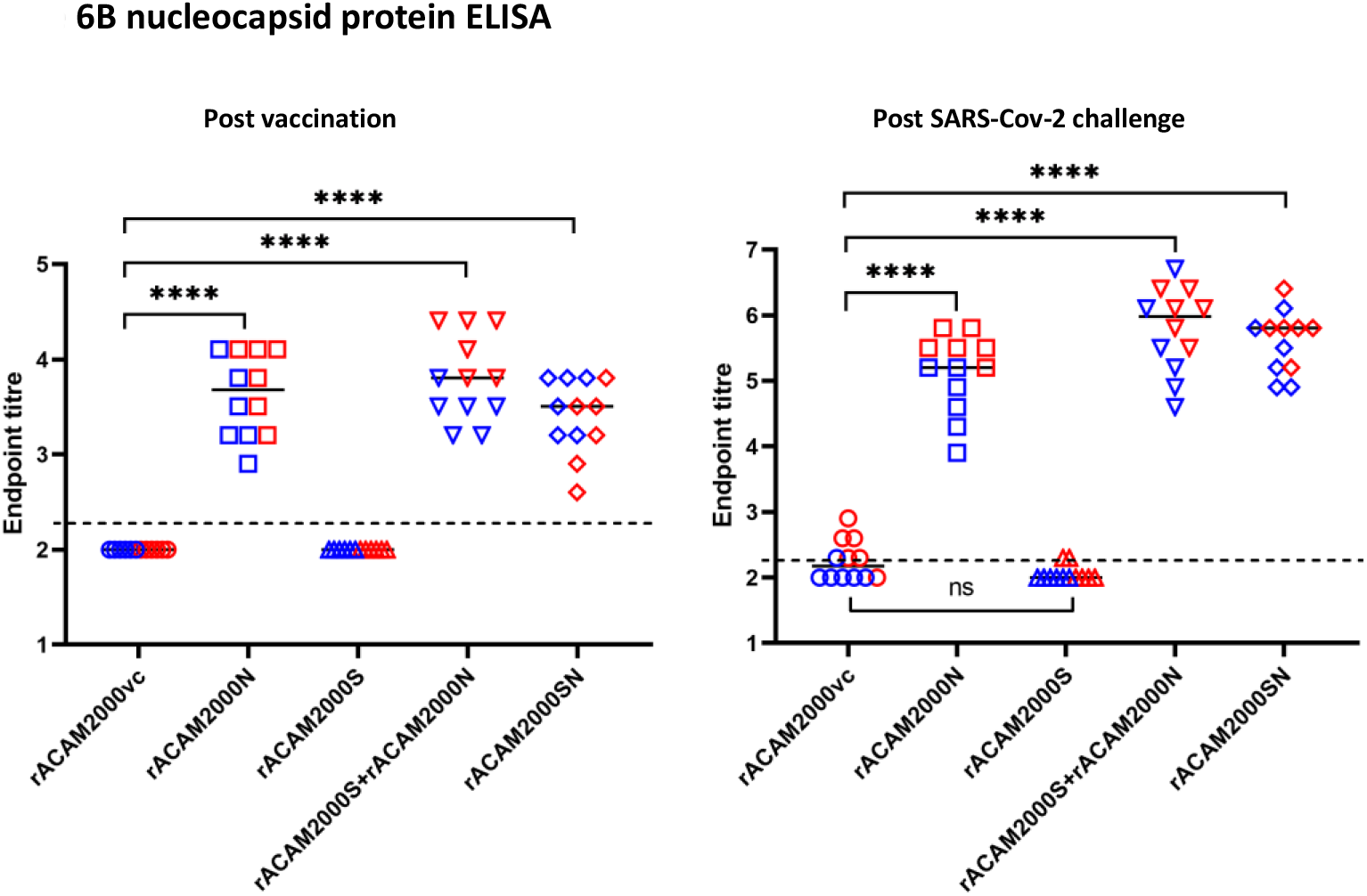
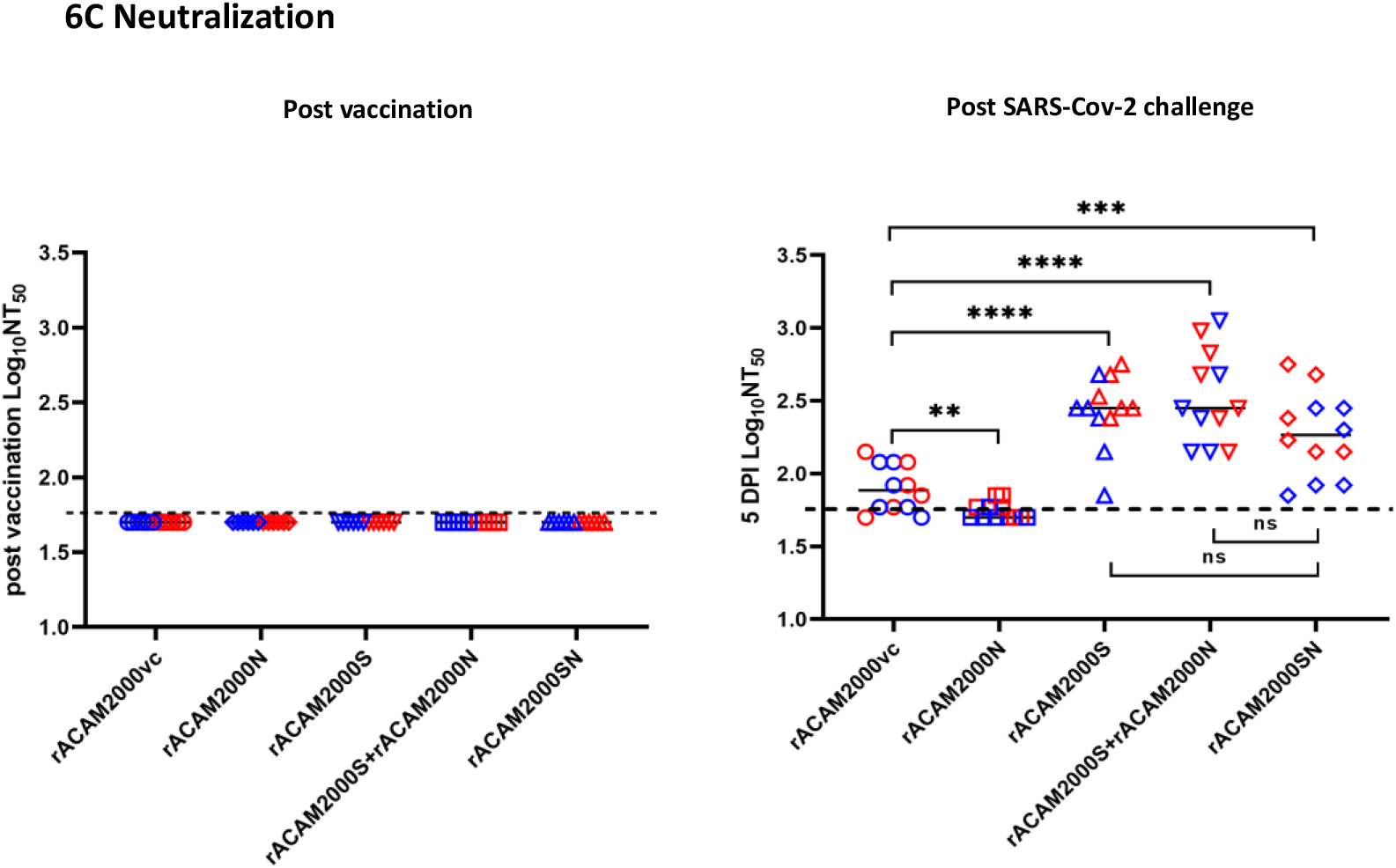
Antibody response at the 21 days post vaccination and 5 days post SARS-CoV-2 challenge. A: Antibody binding the S protein ELISA. The endpoint titers were determined as log_10_ values of the reciprocal of highest serum dilution at which the OD450 was at least two fold higher than the pre-vaccination sera. B: Antibody binding to the N protein ELISA. The endpoint titers were determined the same as in A. C: Neutralizing antibody in the serum. The titers were determined as log_10_ values of the reciprocal of serum dilution at which 50% infectivity of 100 plaque forming units (pfu) of SARS-CoV-2 were neutralized. The dotted line indicate the limit of detection. The circle shape represents rACAM2000vc, square for rACAM2000N, triangle for rACAM2000S, upside-down triangle for rACAM2000S+rACAM2000N, and diamond for rACAM2000SN; red-colored shape for female and blue-colored shape for male hamsters. The P values were calculated with two-tailed Mann-Whitney u test using Graphpad Prism 8.0 package, ****:p<0.0001; ***:p<0.001; **:p<0.01; *:p<0.1.

When the sera from male and female hamsters were further comparatively analysed, it was found that female hamsters vaccinated with the rACAM2000vc and rACAM2000N developed significantly higher levels of antibody than the male in the N ELISA 5 days post SARS-CoV-2 challenge (supplemental Figure 5). A similar trend was also observed in other groups immunized with rACAM2000N in the N ELISA and the neutralization assay following SARS-CoV-2 challenge, albeit statistically insignificant (supplemental Figure 5 and 6). No such variation was observed between male and female serum samples in the S ELISA (supplemental Figure 7).

### Immunization with the rACAM2000 COVID-19 vaccine candidates reduced neutrophil-to-lymphocyte ratio (NLR)

The whole blood collected at 5 days post SARS-CoV-2 challenge was analyzed for various hematological parameters. The reference values were the average between males and females from normal hamsters provided by Charles River Laboratories and included in Figure 7 for comparison^42^ (Figure 7). The total white blood cell counts (WBC), although slightly higher than the reference value, were similar among the vaccinated groups and the vector control hamsters following SARS-CoV-2 challenge (Figure 7A). However, the median lymphocyte numbers from vaccinated hamsters were generally higher than the vector controls, although the differences between some vaccinated groups (rACAM200N and rACAM2000SN) and the rACAM2000vc group were not statistically significant (Figure 7B). The opposite pattern was observed in the neutrophil counts (Figure 7C). Thus, the percentage of lymphocytes in the total WBC of the hamsters immunized with rACAM2000 expressing SARS-CoV-2 antigens was higher than the vector controls, while the percentage of neutrophils was lower (Figure 7D, 7E). As a result, the median of the NLR of the vaccinated animals was lower than the control hamsters, albeit the difference between the NLR from the hamsters vaccinated with rACAM2000N and the rACAM2000vc group was not statistically significant (Figure 7F). No significant difference in the hematology parameters was observed between male and female hamsters (supplemental Figure 8).

**Figure 7.**
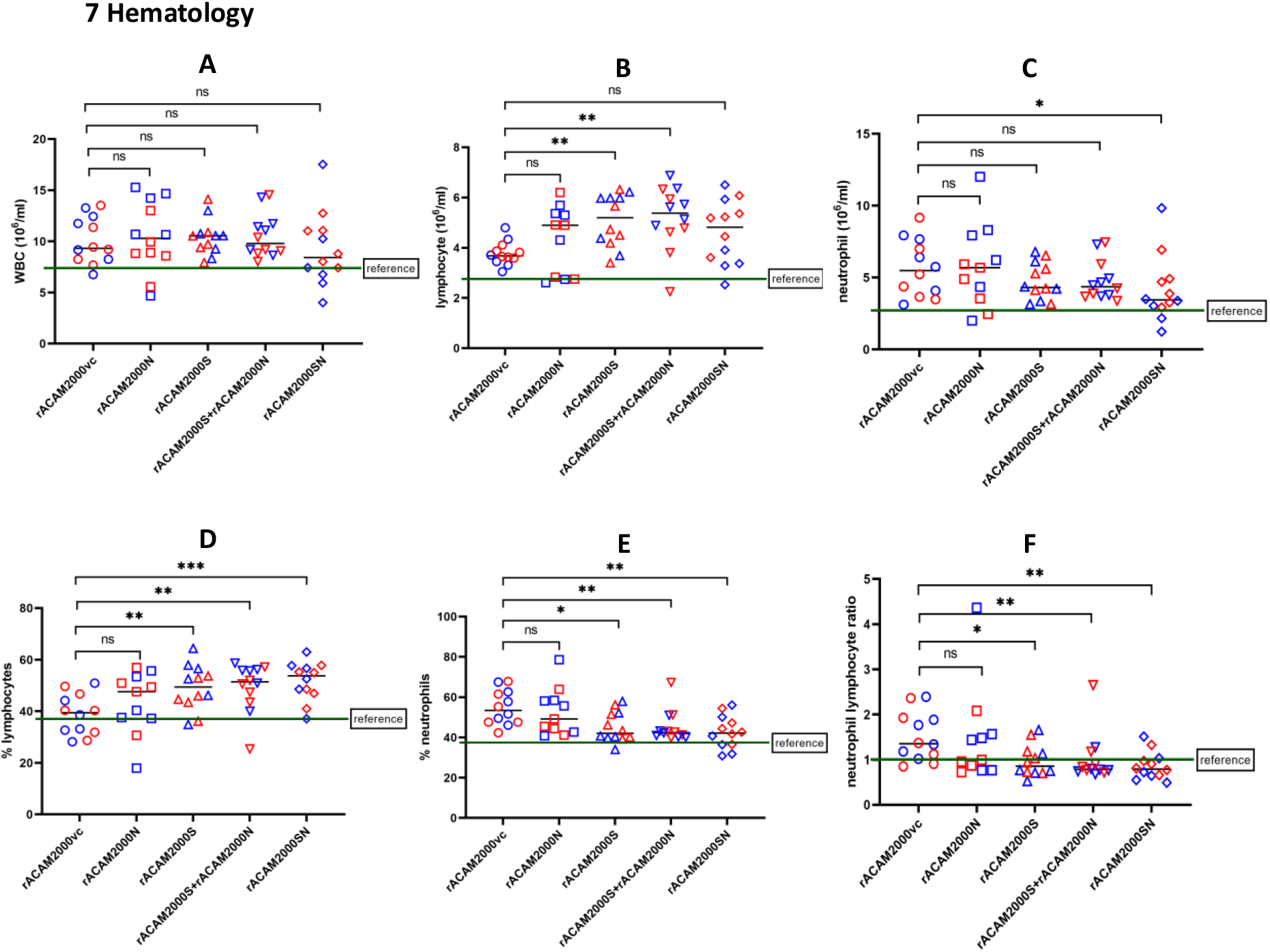
Hematology of hamsters vaccinated and challenged with SARS-CoV-2 at 5DPI. A: total white blood cells (WBC) count; B: lymphocyte count; C: neutrophil count; D: lymphocyte percentage in the total WBC; E: neutrophil percentage in the total WBC; F: neutrophil-to-lymphocyte ratio. The circle shape represents rACAM2000vc, square for rACAM2000N, triangle for rACAM2000S, upside-down triangle for rACAM2000S+rACAM2000N, and diamond for rACAM2000SN; red-colored shape for female and blue-colored shape for male hamsters. The P values were calculated with two-tailed Mann-Whitney u test using Graphpad Prism 8.0 package, ****:p<0.0001; ***:p<0.001; **:p<0.01; *:p<0.1.

### Histopathology

Histopathological analysis of the lung was performed 5 days post SARS-CoV-2 challenge. Representative images of hematoxylin and eosin stained, formalin-fixed, paraffin–embedded tissues from male and female hamsters are shown at 2X and 20X magnifications in Figure 8A. Gross lesions were evident in all lung tissue with varying severity of pulmonary pathology. Generally, the rACAM2000vc vaccinated animals showed more severe interstitial pneumonia with bronchiolitis, necrosis, edema and hemorrhage regardless of the gender of the animals in comparison to the animals vaccinated with rACAM2000 expressing the S and/or N proteins. Pulmonary pathology from lungs of animals vaccinated with rACAM2000N showed less severe infiltration of leucocytes than in rACAM2000vc vaccinated animals, while those vaccinated with other constructs showed some degree of leucocyte infiltration into airway lumen. The extent of interstitial pneumonia across each lung lobe section was estimated in comparison to non-infected control hamsters (Figure 8B). The overall percentage of the lung tissue with interstitial pneumonia was less in the hamsters vaccinated with rACAM2000 expressing the S and/or N proteins than the rACAM2000vc group, albeit the difference in the case of some constructs, e.g. rACAM2000N and rACAM2000SN was not statistically significant. It is interesting to note that the individual variation in the percentage of interstitial pneumonia lung tissue among hamsters vaccinated with the S and/or N constructs (ranging from approximately 25% to 100%) was greater than the rACAM2000vc group (ranging from 70% to 100%) (Figure 8B).

**Figure 8.**
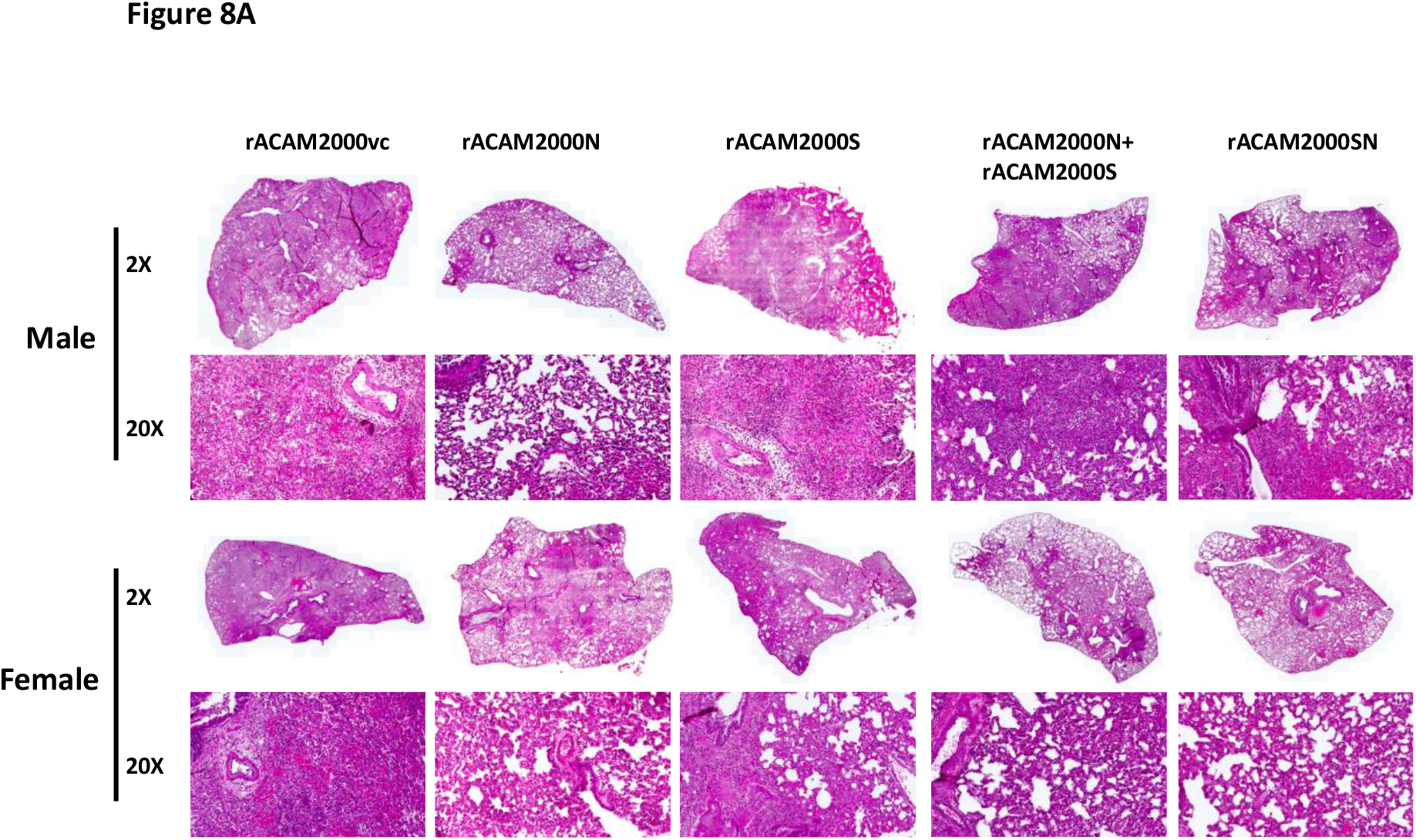
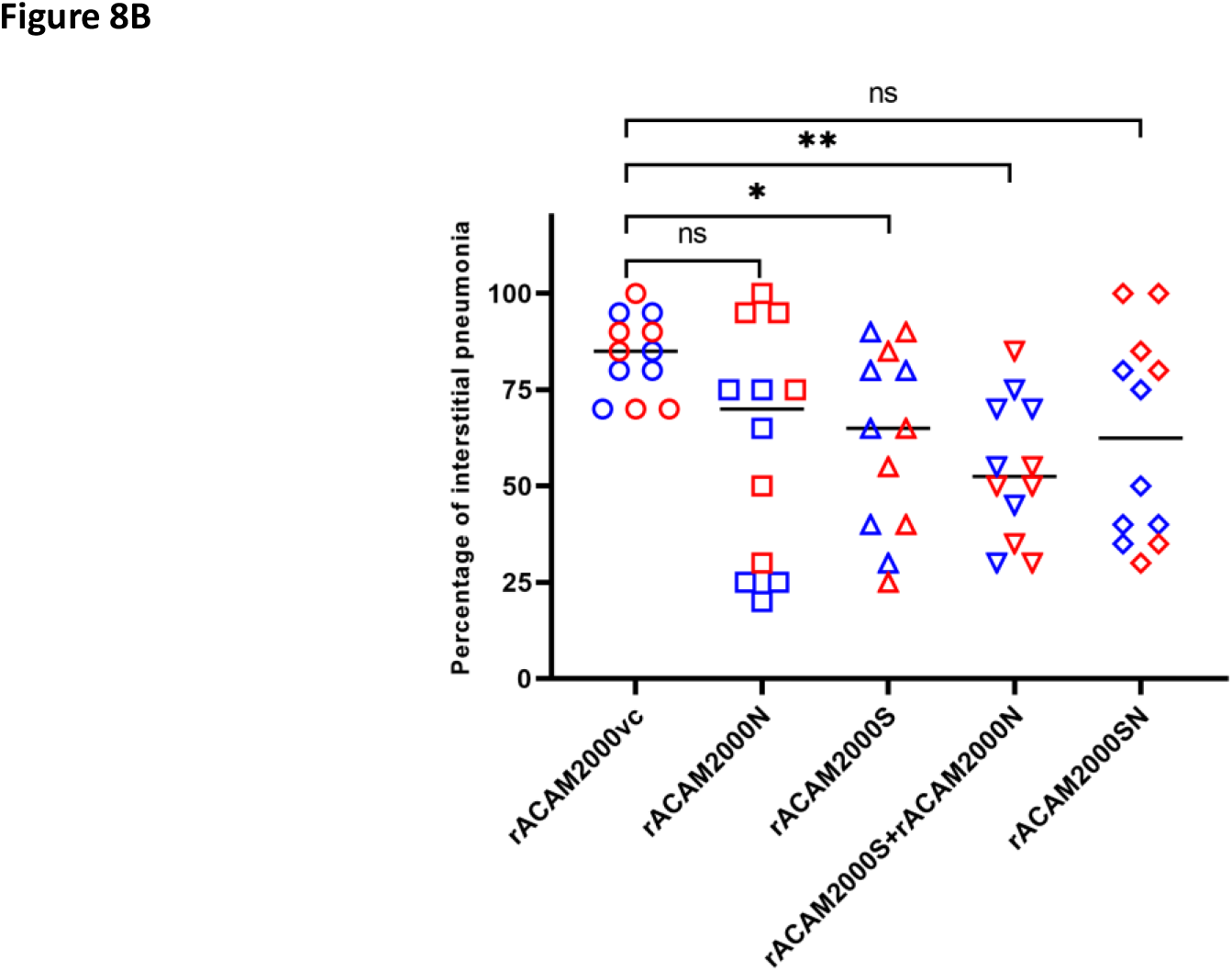
Histopathological analysis. A: Representative images of hematoxylin and eosin stained, formalin-fixed, paraffin–embedded tissues from male and female hamsters at 2X and 20X magnifications. B: Percentage of marked interstitial pneumonia from each hamster lung lobes.

## DISCUSSION

Most currently used COVID-19 vaccines are based on the S protein of the original SARS-Cov-2 Wuhan isolate and they have been effective in preventing the early waves of the pandemic. However, with continuous spreading of the virus, SARS-CoV-2 variants which can cause breakthrough infections from the immunity induced by the vaccines are rapidly emerging^33^. Although the administration of a third dose of the current vaccine (i.e. BNT162b2 mRNA vaccine) has been shown to be effective^43, 44^, new vaccines, which are capable of inducing long-lasting protection against a broad range of SARS-CoV-2 variants, are needed to efficiently combat the ongoing COVID-19 pandemic. In this study, we performed an initial investigation on the protective efficacy of the recombinant COVID-19 vaccine candidates expressing both the S and N antigens based a novel VACV ACAM2000 platform.

MVA, the most popular VACV vector for recombinant vaccines^10^, has been used as one of the major viral vectors for development of COVID-19 vaccine candidates. Several studies have reported that a recombinant MVA expressing the S protein induced protective immune responses in transgenic mice expressing human ACE2 and in non human primate (NHP) model^11–14^. More recently, a recombinant MVA expressing both the S and N proteins was shown to induce neutralizing antibody and cell-mediated immune responses^16^ and was protective against SARS-CoV-2 challenge in hamsters and NHPs^17^. Although MVA is well known for its safety record and capacity to express foreign proteins, Its immunogenicity is lower than the first generation of VACV^27, 45–47^. Moreover, for the application as a recombinant vaccine vector, it has been shown that MVA vectored vaccine candidates tend to work well as a booster rather than primary vaccine in a heterologous prime-boost regimen^48^. Thus, there is a need to develop alternative VACV vectors to improve immunogenicity.

Previously, we developed a fast and efficient method to construct recombinant VACV using the platform based on deletion/exchange of the host range genes E3L and K3L^30^. Since VACV E3L is a virulence gene and the protein it encodes is a potent inhibitor of host innate antiviral immunity, deletion of the E3L gene renders the virus attenuated and more immunogenic than wild-type VACVs^23, 25, 26^. In addition, exchange of the authentic VACV K3L gene with a poxvirus ortholog TATV037 makes the recombinant virus replication-competent in human cells, and thus more robust at expressing a target antigen protein^32^. Recently, we used this platform to construct a recombinant RSV vaccine candidate based on VACV WR strain (a laboratory strain VACV). It was found that the recombinant virus induced comparable protective immune response to the MVA based recombinant in cotton rats and was avirulent in mice^31^. In this study, we constructed recombinant COVID-19 vaccine candidates based on the FDA-approved VACV ACAM2000, a plaque-purified derivative from a widely used VACV strain NYCBH in the eradication of smallpox. We anticipate that the modification of rACAM2000, (i. e. deletion of the host range gene E3L and exchange of the K3L) would significantly reduce the risk of adverse effects of the virus in humans, enhance the immunogenicity, and retain its capacity for expressing a heterogeneous protein for vaccine purposes.

Increasing evidence has demonstrated that variants of SARS-CoV-2 were partially resistant to the neutralization induced by the currently deployed S protein based vaccines^4–8^. Since the N protein is an abundantly expressed protein and more conserved among various SARS-like viruses than the S protein^49^, it is conceivable that combining the S and the N proteins in the vaccine design would likely provide improved protection against the emerging variants. Conserved T-cell epitopes in the N protein have been identified among SARS-CoV-2 variants and these epitopes may induce cross-protective immune responses against a wide range of emerging variants of SARS-CoV-2^50, 51^. Several studies have demonstrated that the N protein vaccine candidates based on different platforms induced protection against SARS-CoV-2 infection^17, 52–56^. In this study, we constructed three novel rACAM2000 based COVID-19 vaccine candidates: rACAM2000S, rACAM2000N and rACAM2000SN. Their protective efficacy was compared against SARS-CoV-2 challenge in a hamster model.

Hamsters vaccinated with a single dose of constructs expressing both the S and N antigens (rACAM2000S+rACAM2000N or rACAM2000SN) were significantly better protected than hamsters vaccinated with the rACAM2000S or rACAM2000N (Figure 3B, 3C). Thus, our study clearly demonstrates that the immune responses induced by the N protein contribute to the overall protective efficacy. A recombinant COVID-19 vaccine candidate based on MVA expressing both the S and N proteins was shown to protect hamsters against SARS-CoV-2 infection after two doses of vaccination^17^. It would be valuable for the future design of recombinant VACV based COVID-19 vaccines to compare the protective efficacies of the recombinant MVA^17^ and the rACAM2000 (described in this study) vaccine candidates expressing both the S and N proteins. It is interesting to note that the effect from immunization with rACAM2000N on the viral load in the distal tissues, e.g. liver and small intestine, was more significant than the proximal respiratory tissues, such as nasal turbinate and lung. This observation suggests that the rACAM2000N induced immune responses limited the virus dissemination from the primary infection site (i.e. lung). Further studies are required to elucidate the link between the development of clinical disease and the virus dissemination in the hamster model. A similar finding was also reported in a mouse model, in that SARS-CoV-2 viral load in the brain of K18-hACE2 mice immunized with a combination of the S and N expressed from an Adv5 platform was lower than the mice immunized with the S only construct following i.n. challenge with SARS-CoV-2 virus, while there was no significant difference in the viral load in the lung^54^.

Neutralizing antibody is a major indicator of the protective immune response induced by S protein-based COVID-19 vaccines^57^. In sera collected post-vaccination, but prior to the SARS-CoV-2 challenge, none of the vaccinated hamsters developed detectable neutralizing antibody. At 5 days post SARS-CoV-2 challenge, however, the neutralizing antibody titre in the hamsters vaccinated with the recombinants expressing the S protein (rACAM2000S, S+N and SN) was significantly higher than the vector control (rACAM200vc). Therefore, rACAM2000 expressing the S protein (rACAM2000S and rACAM2000SN) induced effective memory B cell response responses for the S protein, which were boosted following SARS-COV-2 infection to produce significant level of neutralizing antibody. Unexpectedly, the neutralizing antibody titres in hamsters vaccinated with rACAM2000N were lower than rACAM2000vc control hamsters following SARS-CoV-2 challenge. In fact, 7 out of 12 hamsters in the rACAM2000N group did not show detectable levels of neutralizing antibody post SARS-CoV-2 challenge, compared to 2 out 12 hamsters in the rACAM2000vc group (Figure 6C). The similar antibody profile was also demonstrated by ELISA. Therefore, the protection induced by rACAM2000N was the effect of cell-mediated immune responses. It is possible that vaccination with rACAM2000N induced a strong Th1 immune response that may have resulted in a weaker anti-spike Ab response. It is interesting to note that hamsters vaccinated with rACAM2000SN also developed lower S antibody levels than the hamsters immunized with the rACAM2000S+rACAM2000N and rACAM2000S (Figure 6A). This could be due to the combined effects of the antigen presentation and/or an unknown biological function of the N protein. The S and N antigens expressed from rACAM2000SN were processed and presented from the same cells, while the two antigens were likely to be expressed, processed and presented from different cells in hamsters immunized with rACAM2000S+rACAM2000N since the two recombinants were injected separately in the right or left quadriceps muscles. Further studies with an animal model, such as a mouse model, which has more readily available reagents to analyze various immune responses, should help to elucidate the protective immune responses and the factors involved in the lower antibody responses associated with the constructs expressing SARS-CoV-2 N protein.

It is well documented that males and females respond to various viral infections and vaccination differently at levels of both innate and adaptive immune responses^58–60^. Among COVID-19 patients, increasing evidences have demonstrated that men tend to develop more severe clinical manifestations^61–65^. To our knowledge, however, it is unknown if the currently used COVID-19 vaccines induce different protection between men and women. In this study, we found that rACAM2000 expressing SARS-CoV-2 N protein protect male and female hamsters differently, in that the vaccinated female hamsters recovered significantly quicker than the males following SARS-CoV-2 challenge. Further investigation of this observation should aid in an improved design of vaccines and therapeutic interventions for COVID-19.

Increased NLR has been shown to be associated with severe cases of COVID-19^66–69^. It is likely that the increased NLR may enhance the severity of COVID-19 by inducing unbalanced immune responses, such as hyper inflammation^70, 71^. In this study, we found that the protection induced by rACAM2000s expressing the S and N proteins was associated with reduced NLR in hamsters following SARS-CoV-2 challenge. To our knowledge, this is the first observation that immunization with a COVID-19 vaccine candidate reduced the NLR. However, it should be noted that the disease caused by SARS-CoV-2 in hamsters is not the same as human COVID-19. Further studies with a different animal model, particularly those with more reagents available for various immunological assays (e.g. a mouse model) should provide more information on the link between immunization, reduced NLR and disease development.

In summary, we evaluated the protective efficacy of three recombinant COVID-19 vaccine candidates expressing SARS-CoV-2 S and N proteins individually or in combination using a novel VACV rACAM2000 backbone. A single dose of vaccination with the rACAM2000 expressing both the S and the N antigens induced significantly better protection against SARS-CoV-2 infection in hamsters than the construct expressing individual S or N proteins. Interestingly, our data showed that vaccination with the rACAM2000N decreased antibody responses and induced sex-dependent protection in hamsters following SARS-CoV-2 challenge. Further investigation to understand the mechanism related to this observation is important for application of the N protein in vaccine design. Studies are underway to investigate if the rACAM2000 based COVID-19 vaccine candidate will induce long lasting protective immunity against emerging SARS-CoV-2 variants of concern.

## Acknowledges

This work was funded by the Public Health Agency of Canada. We thank Carissa Embury-Hyatt of Canadian Food Inspection Agency for histopathological analysis of the preliminary hamster study.

## Author contributions

The overall study was conceived of and designed by J.X.C… Y.D. and J. L. constructed the recombinant shuttle vectors. J.X.C. prepared all the recombinant VACV ACAM2000 viruses. J.L., C.L, and K.L. performed Western blotting and sequence analysis to confirm the recombinant viruses. J.X.C., B.W. and D.K. designed the animal experiment and B.W. and K.D. prepared animal user’s document. Y.D., K.T. and K.D. performed all the animal procedures and Y.D. coordinated all the animal experiments. R.V. and N.T. performed blood chemistry and hematology analysis. Y.D. performed micro-neutralization experiment. D.H., J.L., and C.L. performed ELISA assays. M.C., L.L., and D.B. produced purified S and N proteins used for ELISA. Y.D. performed TCID_50_ assay and J.L. performed qRT-PCR assay. S.B., K.F., and B.S. performed histopathological analysis. Data were interpreted by J.X.C., X.L., D.K., D.S., S.B. and B.W… The draft manuscript was written by J.X.C., and all the authors contributed to the editing of the paper and the final version of the paper.

---

**Supplemental Figure 1.**
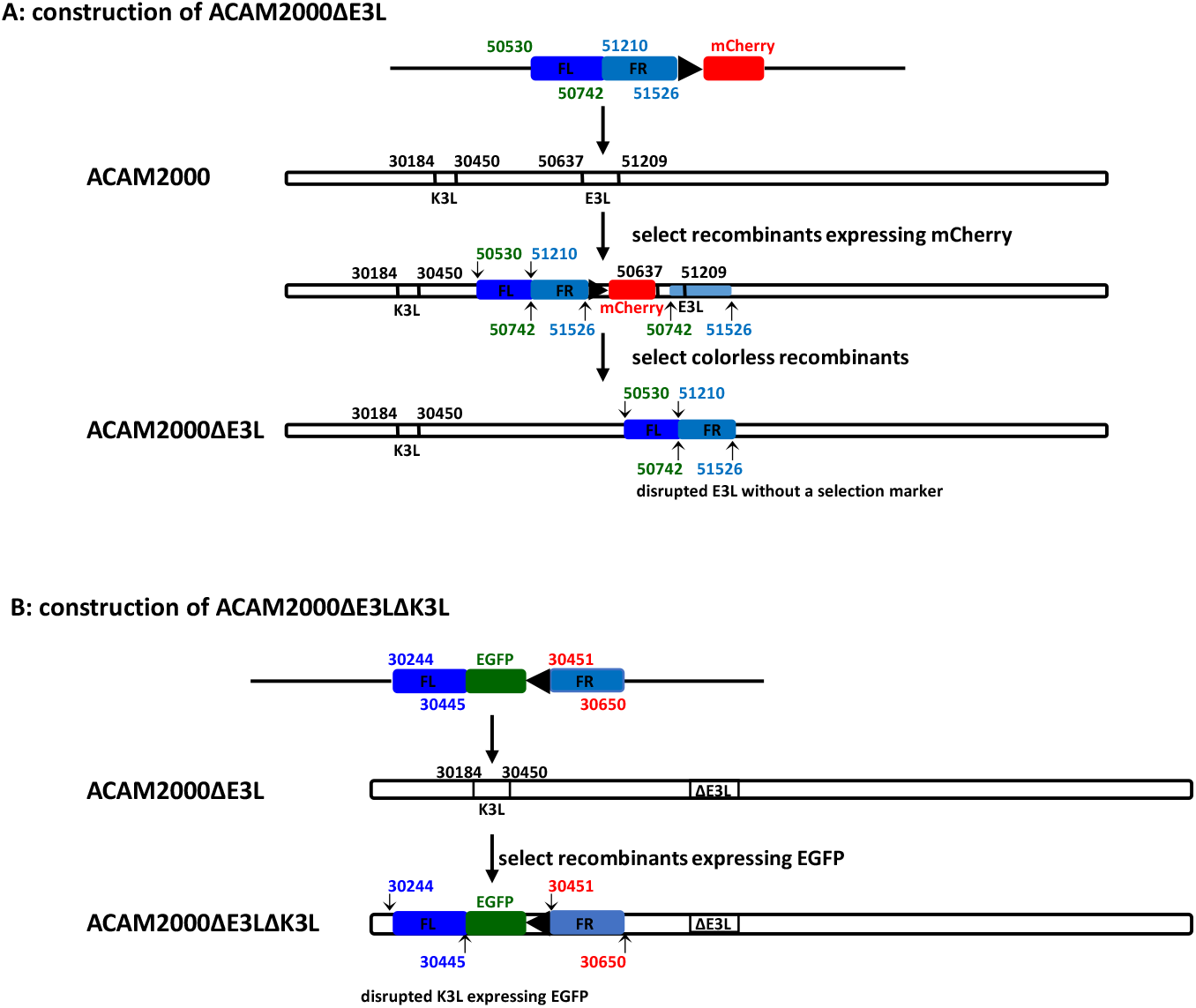
Construction of E3L and K3L deletion mutant of VACV ACAM2000. A: Schematic illustration of construction of the E3L deletion mutant of VACV strain ACAM2000 (ACAM2000ΔE3L). The numbers on the recombinant shuttle vector and the virus genome correspond to the base pair number in the genome of ACAM2000 (accession number AY313847). The transient selection marker mCherry gene is driven by the vaccine virus late promoter p11. The final ACAM2000ΔE3L virus has the E3L gene disrupted without the transient selection marker mCherry as colorless plaques were selected during the virus purification. B: Schematic illustration of construction of the E3L and K3L double deletion mutant (ACAM2000ΔE3LΔK3L). The numbers on the recombinant shuttle vector and the genome of ACAM2000ΔE3L correspond to the base pair number in the genome of ACAM2000 (accession number AY313847). The selection marker EGFP gene is driven by the vaccinia late promoter p11. The final ACAM2000ΔE3LΔK3L has both E3L and K3L genes disrupted and with an insertion of EGFP in the locus of the K3L gene.

**Supplemental Figure 2.**
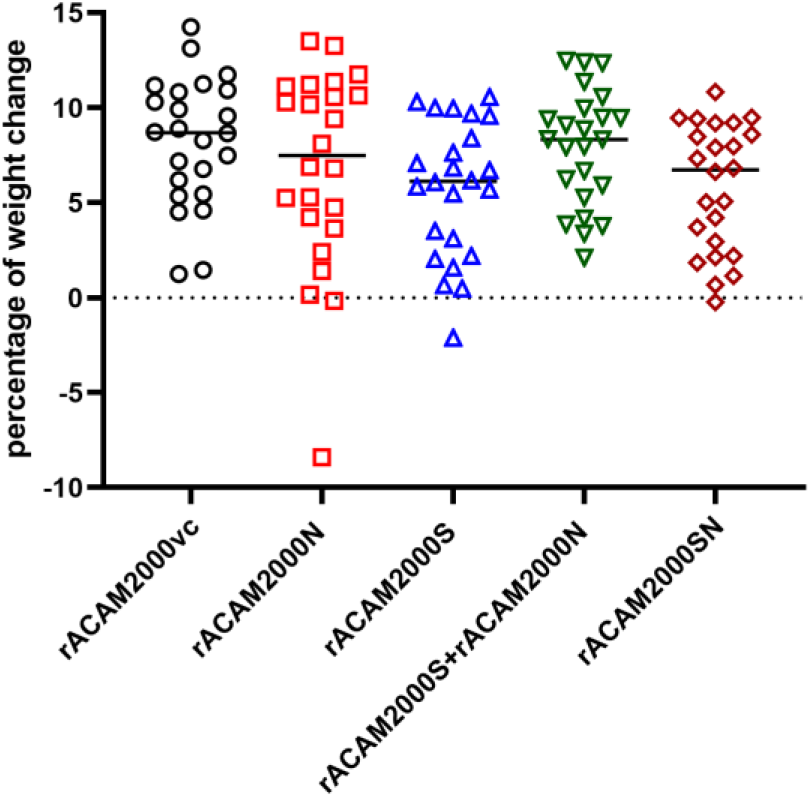
Weight change 7 days post vaccination. Percentage of the weight change of all hamsters 7 days post the vaccination in comparison to days 0.

**Supplemental Figure 3.**
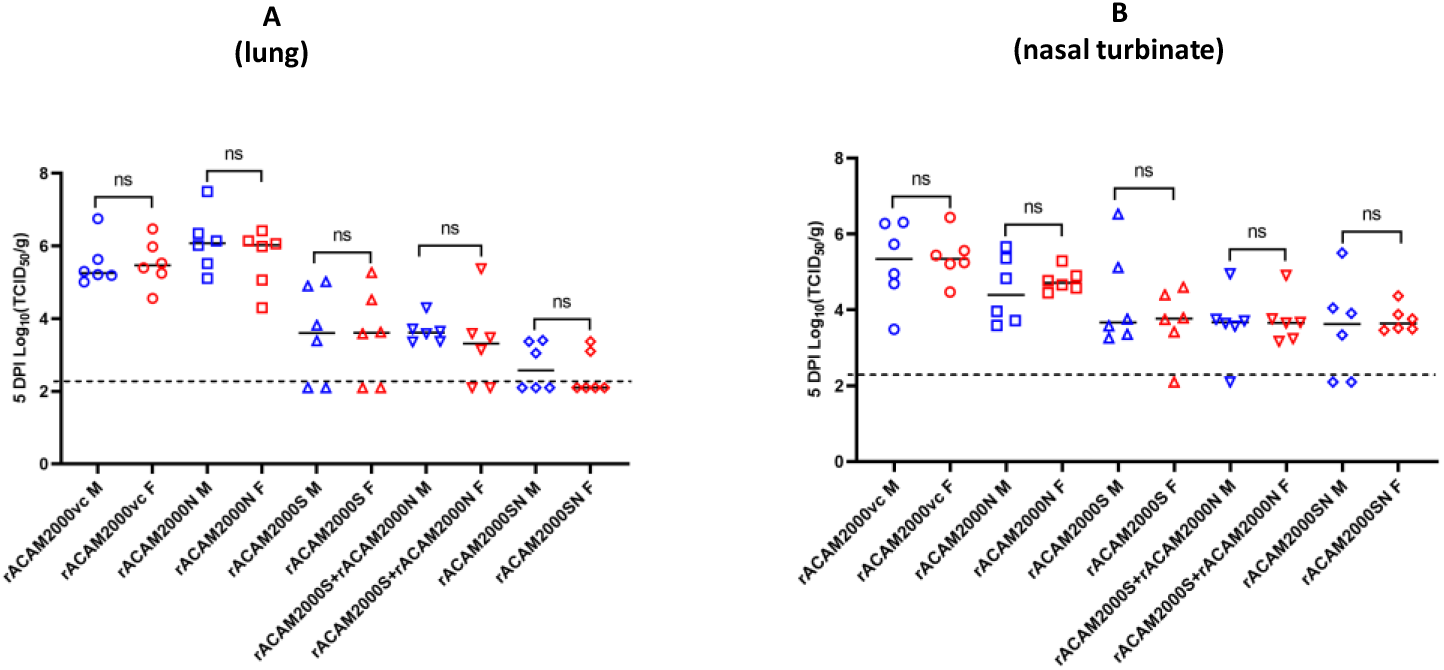
Comparison of infectious SARS-Cov-2 loads between male and female hamsters. The infectious SARS-CoV-2 virus loads determined by TCID_50_ assay shown in Figure 4 were further analyzed by comparing the virus loads of male and female hasmters. M: male; F: female.

**Supplemental Figure 4.**
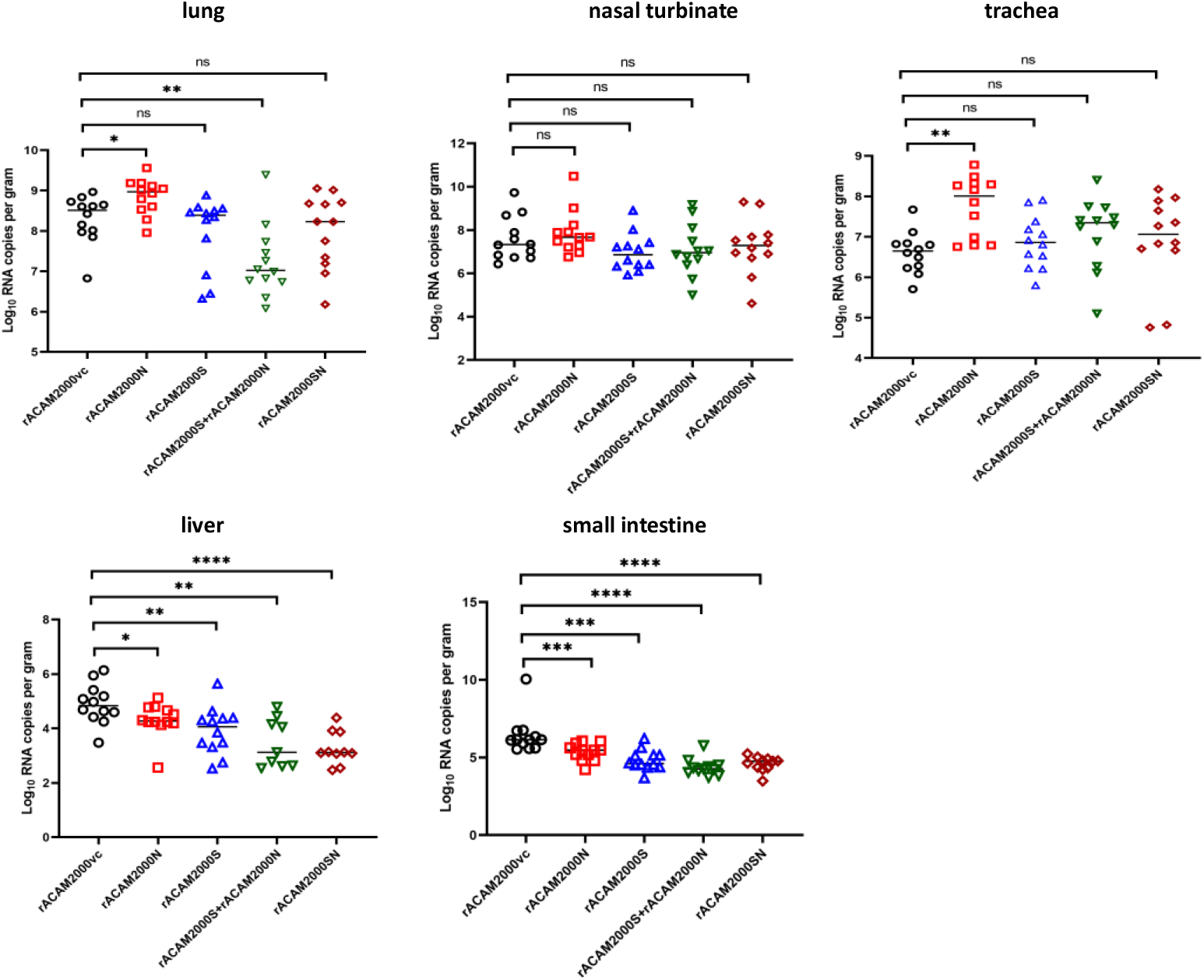
SARS-Cov-2 genomic RNA copy numbers determined by qRT-PCR with the N primers. A: The RNA copy numbers (per gram of tissue) in the lung. B: The RNA copy numbers in the nasal turbinate. C: The RNA copy numbers in the trachea. D: The RNA copy numbers in the liver. E: The RNA copy numbers in the small intestine.

**Supplemental Figure 5.**
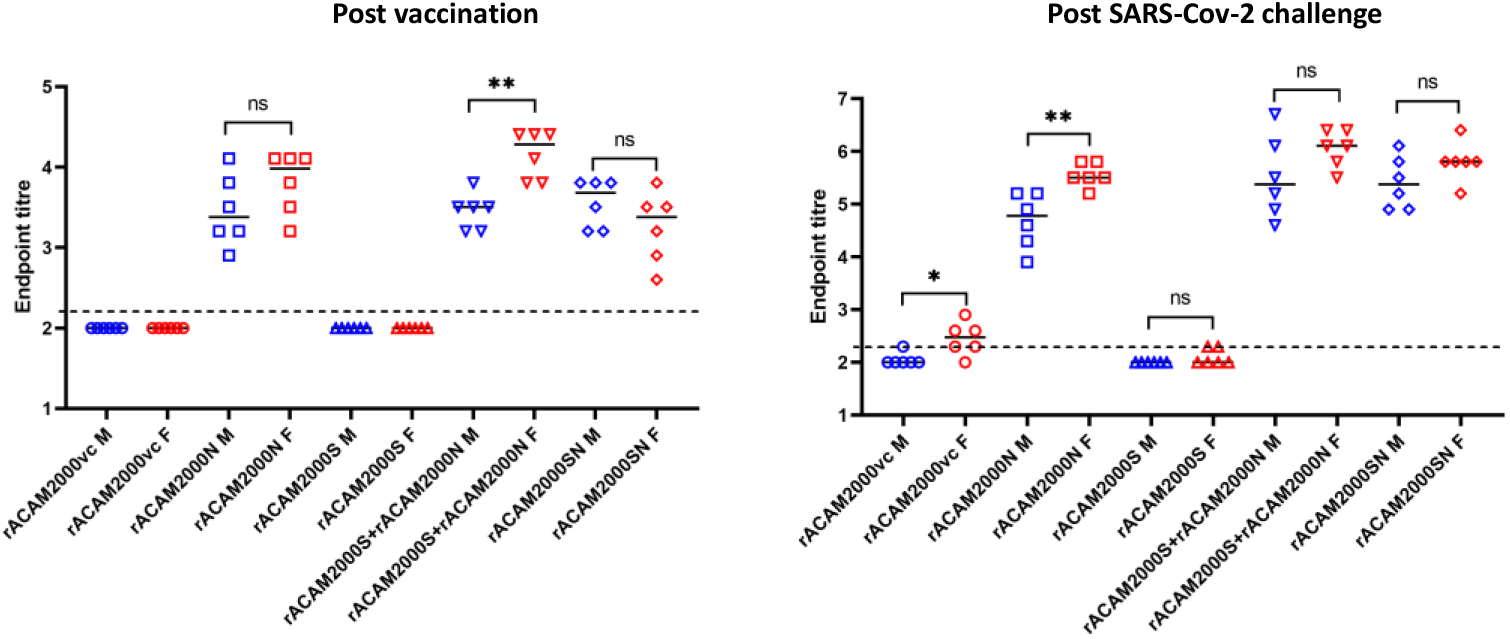
Comparison of the antibody titers against N protein between male and female hamsters using ELISA. Antibody levels against the N protein post vaccination and SARS-CoV-2 challenge were compared between male and female hamsters. The antibody titer was determined as in Figure 6B. M: male; F: female.

**Supplemental Figure 6.**
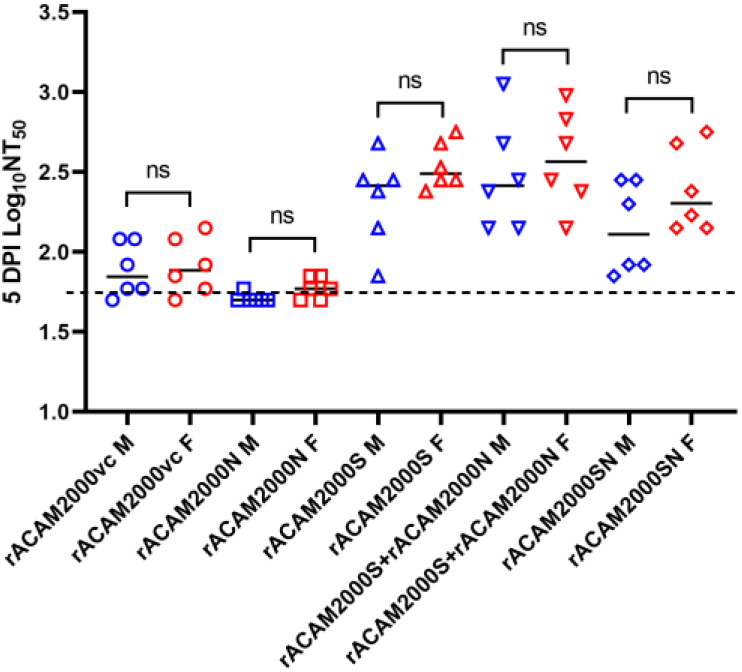
Comparison of neutralizing antibody titers between male and female hamsters. The neutralizing antibody titers were compared between male and female hamsters as shown in Figure 6C. M: male; F: female.

**Supplement Figure 7.**
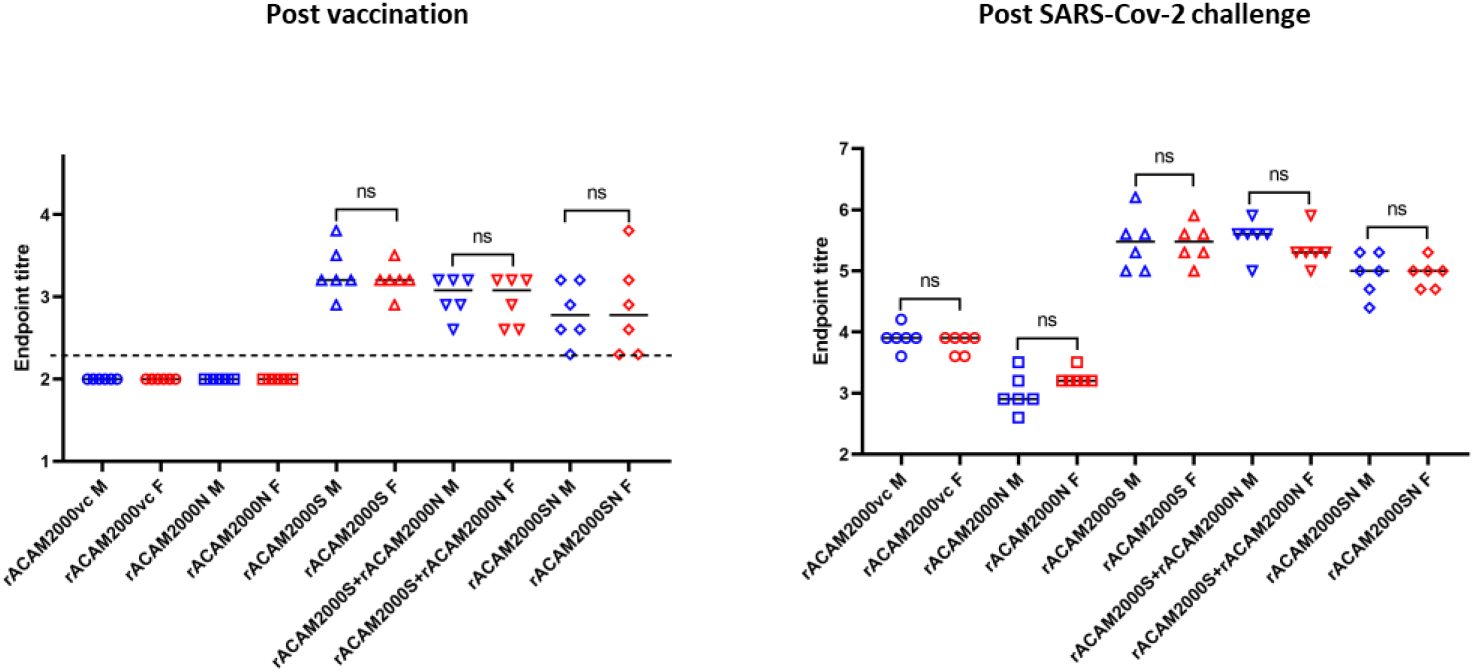
Comparison of the antibody titers against S protein between male and female hamsters using ELISA. Antibody levels against the S protein post vaccination and SARS-CoV-2 challenge were compared between male and female hamsters. The antibody titer was determined as in Figure 6A. M: male; F: female.

**Supplemental figure 8.**
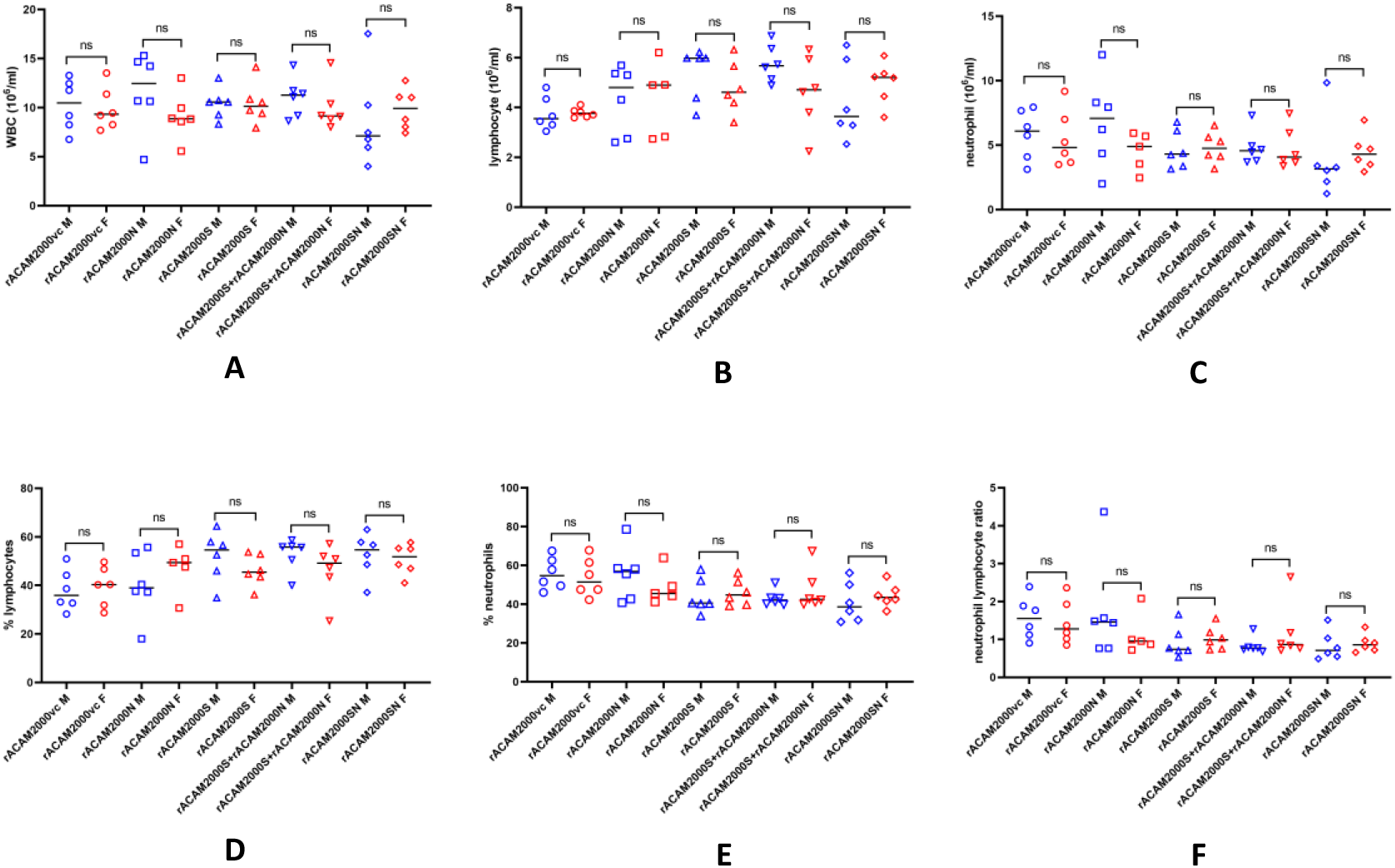
Comparison of hematology between male and female hamsters. The hematological data were from Figure 7; male and female data were compared. . M: male; F: female.

